# Evolutionary origin and structural ligand mimicry by the inserted domain of alpha-integrin proteins

**DOI:** 10.1101/2023.11.05.565221

**Authors:** Jeremy A. Hollis, Matthew C. Chan, Harmit S. Malik, Melody G. Campbell

## Abstract

Heterodimeric integrin proteins transmit signals through conformational changes upon ligand binding between their alpha (α) and beta (β) subunits. Early in chordate evolution, some α subunits acquired an “inserted” (I) domain, which expanded their ligand binding capacity but simultaneously obstructed the ancestral ligand-binding pocket. While this would seemingly impede conventional ligand-mediated integrin activation, it was proposed that the I domain itself could serve both as a ligand replacement and an activation trigger. Here, we provide compelling evidence in support of this longstanding hypothesis using high-resolution cryo-electron microscopy structures of two distinct integrin complexes: the ligand-free and E-cadherin-bound states of the αEβ7 integrin with the I domain, as well as the α4β7 integrin lacking the I domain in both a ligand-free state and bound to MadCAM-1. We trace the evolutionary origin of the I domain to an ancestral collagen-collagen interaction domain. Our analyses illuminate how the I domain intrinsically mimics an extrinsic ligand, enabling integrins to undergo the canonical allosteric cascade of conformational activation and dramatically expanding the range of cellular communication mechanisms in vertebrates.

## Introduction

Protein evolution often proceeds via relatively small incremental mutational steps. In contrast, gain of protein domains can spur dramatic novelty, sometimes at the cost of ancestral functions. One such dramatic domain acquisition happened early in chordate evolution in the integrin cell surface receptor family, which mediates signaling across the cell membrane in a range of biological processes from embryogenesis to T-cell activation. Integrin molecules are heterodimers composed of a single alpha (α) and beta (β) subunit (Supplemental Fig. 1A). The human genome encodes 8 β and 18 α integrin subunits, 9 of which include a 200aa derived “inserted” von Willebrand factor type-A (vWFA) domain (or I domain)^1–4^. Phylogenetic analyses have revealed that the acquisition of the I domain was a monophyletic event in the common ancestor of all *Olfactores*, the clade encompassing urochordates (tunicates) and vertebrates^5^. Ancestrally, α-integrin proteins encode seven repeats, each coding for a single blade of a seven-blade beta-propellor fold. The I domain is placed between the second and third propeller blades^6^ (Supplemental Fig. 1B).

Originally identified in the von Willebrand factor protein^7^, vWFA domains typically adopt a classic Rossmann fold and contain a metal-ion dependent adhesion site (MIDAS) for binding protein ligands in a variety of contexts^8^. Integrin-ligand binding was ancestrally mediated by an interface between α and β subunits. However, the vWFA-family I domain insertion sterically occluded this ligand-binding interface. Instead, the I domain itself acquired ligand binding functions, enabling integrin heterodimers to bind a much wider array of ligand moieties and motifs^9^. Ligand binding in ancestral integrins is further directly linked to the conformational changes required to signal integrin activation (Supplemental Fig. 1C) by a canonical Arg-Gly-Asp (RGD) tripeptide present in many extrinsic ligands^10^. Acquisition of the I domain also potentially blocked the trigger for integrin activation. One resolution to this conundrum emerged from previous structural characterization of isolated I domains, which revealed a conformational change in their C-terminal helix that inserts a conserved glutamate into the ancestral ligand binding pocket of integrin^11^. This finding led to the proposed “internal ligand” model in which the I domain functionally replaces the canonical RGD tripeptide that is required for activation of ancestral, non-I domain integrins^12^.

Despite the elegance and explanatory power of the “internal ligand” model, our current structural understanding of how I domain-triggered integrin activation occurs is still greatly limited. A significant hurdle has been methodology; integrin ectodomains preferentially crystallize in a compact (inactive) state. As a result, to our knowledge the conformational landscape for an I domain-containing integrin has never been observed with the required structural detail to resolve features of ligand binding and integrin activation; additional unknown contacts beyond the conserved glutamate are likely critical^12^. Moreover, the limited biochemical support for the “internal ligand” model was obtained using structures of β2 integrins, which exclusively pair with I-domain containing α-integrin subunits and might possess idiosyncratic adaptations not shared with I domain-less integrin heterodimers. This precludes direct comparisons between previous structures. Given that half of human α-integrins, including the vast majority of those with critical immune function, contain I domains, resolving the molecular details of this evolutionary insertion would provide vital insight into a wide array of vertebrate signaling processes.

We directly tested the intrinsic ligand model using single-particle cryogenic electron microscopy (cryoEM) analyses. Recent advances in cryoEM can resolve sample heterogeneity and determine structural ensembles^13-15^, which allows us to characterize the complicated allosteric relay required for integrin activation.L We directly compare the activation networks of two integrin heterodimers involving β7, one of only two β integrin subunits capable of interacting with both types of α-integrin subunits (Fig 1A). We characterize the α4β7 (or LPAM-1) heterodimer, consisting of the I domain-lacking α4 subunit (or CD49d) in complex with β7, and compare this to αEβ7 integrin heterodimer, consisting of the I domain-containing αE subunit (or CD103) and β7. Both integrins perform critical roles in facilitating T cell homing, activation, and retention at sensitive immune barriers^16,17^, yet molecular details of their ligand binding and activation are largely unknown. These integrins are promising therapeutic candidates for the treatment of Irritable Bowel Diseases^18,19^ where there is scope for improving efficacy^20,21^. Structural insight into the mechanism of activation of the α4β7 and αEβ7 integrins could reveal the molecular contacts mediating β7 integrin activation and ligand engagement, providing the framework needed to develop precisely targeted, next-generation therapeutics for gut mucosal inflammation and autoimmunity. Moreover, structures of αEβ7/CD103 will lead to rationally-designed cancer immunotherapies that exploit the prognostically favorable tissue resident-memory T-cell (T_rm_) phenotype, effectuated by the αEβ7:E-cadherin interaction^22^. In addition to the evolutionary insights we uncover, structures of these β7 integrins have enormous clinical value.

**Figure 1.**
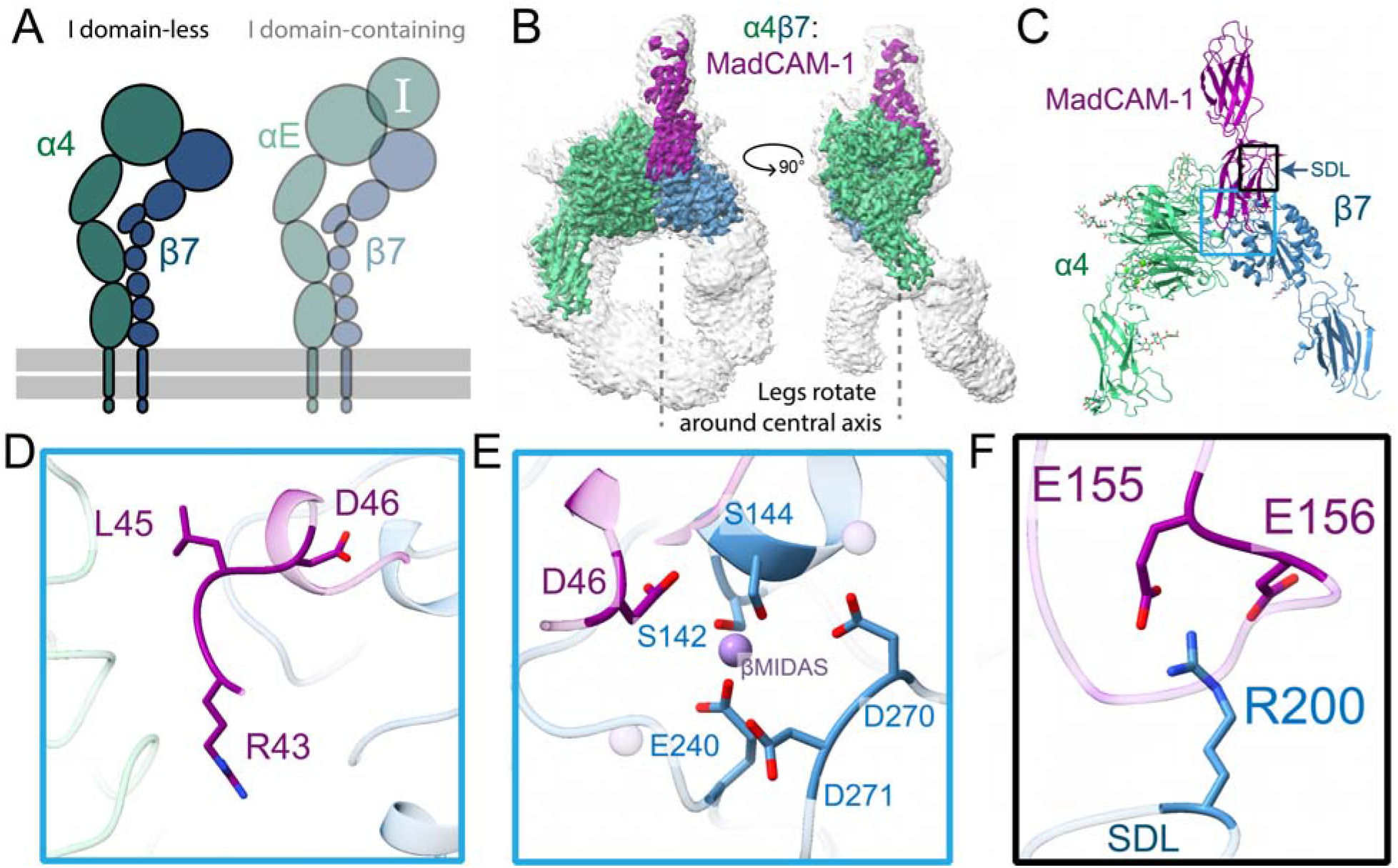
The structure of α4β7 bound to ligand MadCAM-1. **A,** A schematic highlighting showing the focus of this figure on the I domain-lacking integrin α4β7. **B,** CryoEM structure of the integrin α4β7 ectodomain (green/blue) bound to the MadCAM-1 ectodomain (purple). The low threshold, unsharpened global refinement map is shown in gray, and the inset high threshold, sharpened local refinement map is shown in color. The legs of α4 and β7 are not planar; rather they appear to rotate around a vertical axis. **C,** Atomic model of the α4β7:MadCAM-1 complex. Blue box indicates regions of focus in (D-E), black box indicates region of focus in (F). **D,** MadCAM-1 uses an RGLD motif instead of the canonical tripeptide RGD to bridge the α4 and β7 subunits. The RGLD residues span the long axis of the cleft formed between α4 and β7. **E,** MadCAM-1 D46 completes the β7 βMIDAS ion coordination sphere, here modeled as Mn^2+^. **F,** R200 in the β7 specificity-determining loop (SDL) forms essential salt bridges with a flexible loop in MadCAM-1.

### Structural insights into activation of non-I domain integrin **α**4**β**7

We first determined a structure of the non-I α4β7 integrin ectodomain bound to its cognate gut-specific primary ligand MadCAM-1 to 3.1Å resolution (Fig. 1B-C, Supplemental Fig. 2, Supplemental Table 1). This interaction strictly mediates leukocyte homing to gut tissues^23^, yet the molecular details that define its selectivity are largely unresolved. In contrast to the pocket shape of the RGD-binding αV integrin group, the cleft between α4 and β7 forms a long groove running parallel to the subunit binding interface^24^. We find MadCAM-1 fits snugly within this groove, forming interactions with both integrin subunits (Supplemental Fig. 3A). Contacts primarily occur on the first Ig-like MadCAM-1 domain, between the MadCAM-1 FG strands and α4 propeller, and between the MadCAM-1 CD loop and the β7 βI domain (Supplemental Fig. 3A-B). MadCAM-1 bridges the α4 and β7 subunits with an RGLD motif analogous to the canonical RGD motif described for other I domain-less integrins. Within this RGLD motif, R43 and L45 span the long axis of the integrin groove, filling the space between α4 and β7 (Fig. 1D). The D46 residue provides the essential ion-coordination activity within this motif, leading to the open leg integrin conformation characteristic of an active heterodimer (Fig. 1E). In contrast to existing crystal structures of MadCAM-1 alone^25,26^, we find that the MadCAM-1 D beta strand runs antiparallel to the E strand in Ig-like domain 1 when bound to α4β7. This indicates MadCAM-1 undergoes a conformational change upon binding (Supplemental Fig. 3C). The complex is further supported by the MadCAM-1 DE beta ribbon loop in the Ig-like domain 2 contacting the β7 specificity-determining loop (SDL) via mutationally intolerant^27^ hydrogen bonds (Fig. 1F). However, our model represents only one of many likely electrostatic states continuously sampled by this highly negative MadCAM-1 loop.

We observed a high degree of β7 leg flexibility in the open α4β7:MadCAM-1 structure (Supplemental Video 1). We also observed clear density in the lower leg region of the α4 subunit encompassing the calf 1 and calf 2 domains, undescribed in previous work^24^. We find that the “open” α4β7 conformation has a slight rotation around a vertical axis (Fig. 1B). We hypothesized that α4β7 may “untwist” open rather than simply unfold as suggested in prior structures of other integrins. To understand this potential activation motion, we determined an apo α4β7 clasped structure under high calcium buffer conditions previously shown to favor the inactive conformation state^28^. The resolution of the structure is sufficient to fit domains; however, we do not build an atomic model because the reported resolution of 3.2 Å is likely overestimated due to a preferred orientation (Supplemental Fig. 4A, Supplemental Table 1). In this apo structure, the headpiece regions of α4β7 are not bent 90 degrees relative to the lower legs as with the recently described “half-bent” conformation of the epithelial integrin α5β1^29^ nor is it in the established “acute-bent” conformation. Rather, in this new bent conformation, the headpiece is skewed and β7 headpiece region sits below the α4 propeller. The simplest means to transition from the α4β7 compact to extended state would be by a twist about the lower leg region. Indeed, additional 3D flexibility analysis of the α4β7:MadCAM-1 complex supports rotational movement of the integrin legs (Supplemental Video 1). Given our structural data, it is possible that this “twisted” compact orientation facilitates a similar angle of ligand approach or occlusion between α5β1 and α4β7 even though α4β7 binds MadCAM-1 perpendicular to the orientation at which α5β1 binds fibronectin (Supplemental Fig. 4B).

### The inactive **α**E I domain is dynamic

Next, we turned our attention to structural analysis of I domain-containing αEβ7 integrin heterodimers. To define the conformations of αEβ7 and the relative location of its I domain, we performed negative stain electron microscopy (nsEM) on the purified αEβ7 ectodomain (Supplemental Fig. 5). We observed conformations consistent with both a compact state as well as an open state as seen in nsEM with other integrins^30^. Surprisingly, we found that αEβ7 adopts a “half-bent” conformation similar to α5β1^29^, rather than the acutely bent conformation seen in other I domain-containing leukocyte integrin structures^11,32^. Based on phylogenetic analyses of the respective integrins, our findings suggest that the acutely-bent conformation independently evolved twice in vertebrate integrins. Furthermore, based on previously published EM data^30^ and ours, we infer via parsimony that ancestral integrin must have adopted a half-bent compact conformation (Fig. 2). This result implies a unique and recurring selective pressure on the closed integrin conformation, highlighting the importance of understanding the full integrin conformational spectrum in an evolutionary context.

**Figure 2.**
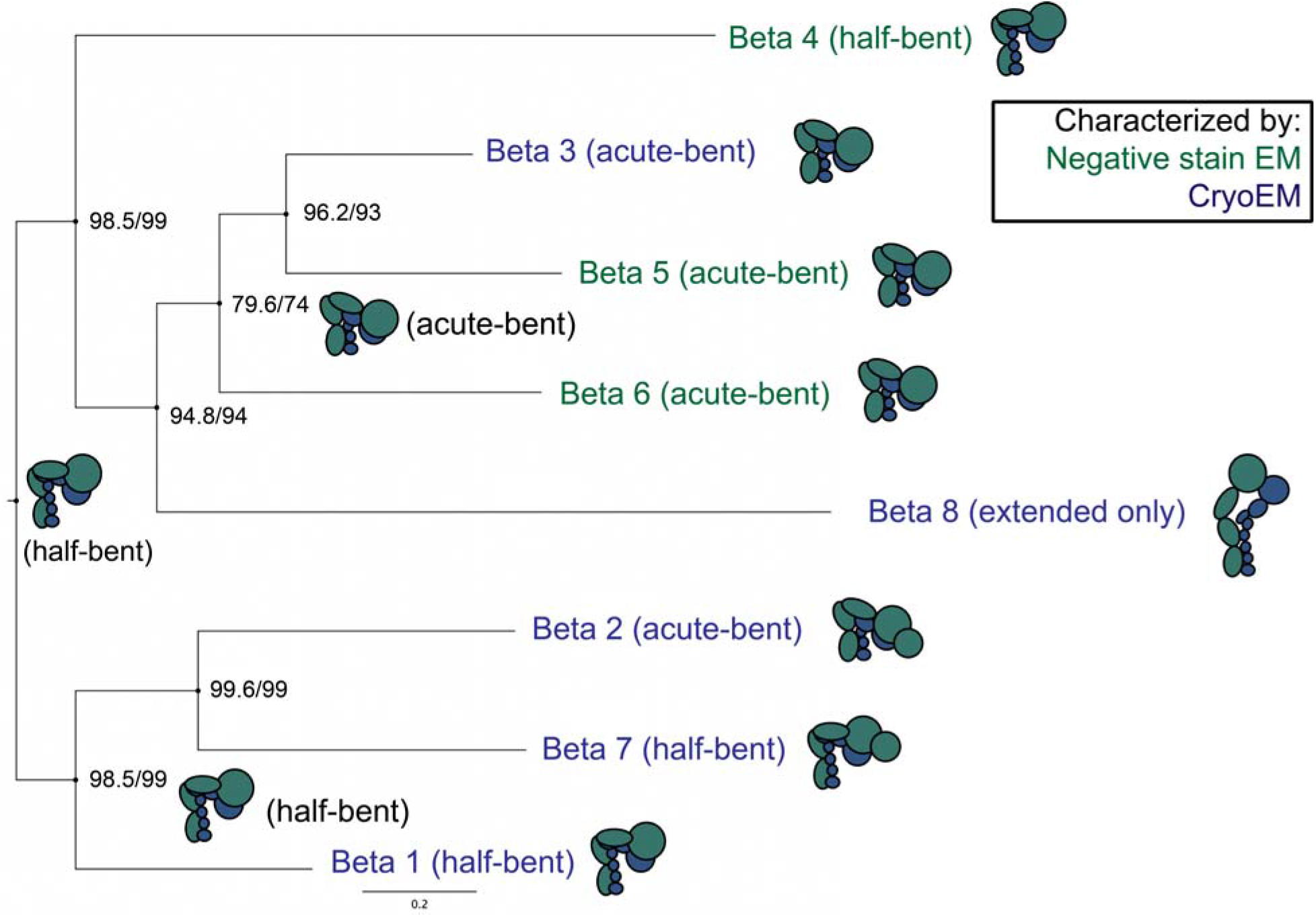
The acute-bent compact integrin state has independently evolved twice. A phylogeny of human beta integrin subunits with compact-state cartoons of integrin dimers was generated with IQ-TREE and annotated with previous ectodomain or full-length integrin structural data. αMβ2^32^, αEβ7 (this study), α5β1^31^, αIIbβ3^53^ and αVβ8^54^ have all been characterized at high resolution. αVβ5^55^, αVβ6^56^ and α6β4^30^ have been characterized at low resolution. The ancestral integrin compact state was most parsimoniously half-bent. Node values are displayed as “ultrafast bootstrap/SH-aLRT”.

To gain more detailed insight into the compact inactive state of the I domain (Fig. 3A), we performed cryoEM studies on the clasped αEβ7 ectodomain under high calcium, or nonactivating, buffer conditions^28^. To facilitate structural analysis, this integrin is bound to a CD103 antibody fragment LF61 that does not influence ligand binding^31^. We resolved the compact structure of the αEβ7 ectodomain to a global resolution of 2.9Å (Fig. 3B, Supplemental Fig. 6). The resolution of our αEβ7 structure was highly anisotropic (Supplemental Fig. 6), but as with α4β7 we could confidently model the αEβ7 headpiece region (Fig. 3C, Supplemental Data Table 1). We identified and modeled five glycosylations on αE, including two on the I domain (Supplemental Fig. 6F). The I domain is positioned above the heterodimer headpiece as expected of an I domain-containing integrin. This position would sterically occlude the ancestral RGD pocket of integrins without an I domain. Even in the compact state, the αE I domain has a more extensive interface with both the β7 and αE subunits than described for β2 leukocyte integrins. We found the αE I domain makes additional contacts with both the β7 SDL and a beta hairpin on the αE beta-propeller. These supports form a “collar” around the αE I domain (Fig. 3D). These additional contacts may contribute to a more stabilized I domain leading to the more resolved structure of αEβ7 I domain than that of αMβ2, which was not resolved in a recent cryoEM density^32^; possibly due to its substantially higher flexibility.

**Figure 3.**
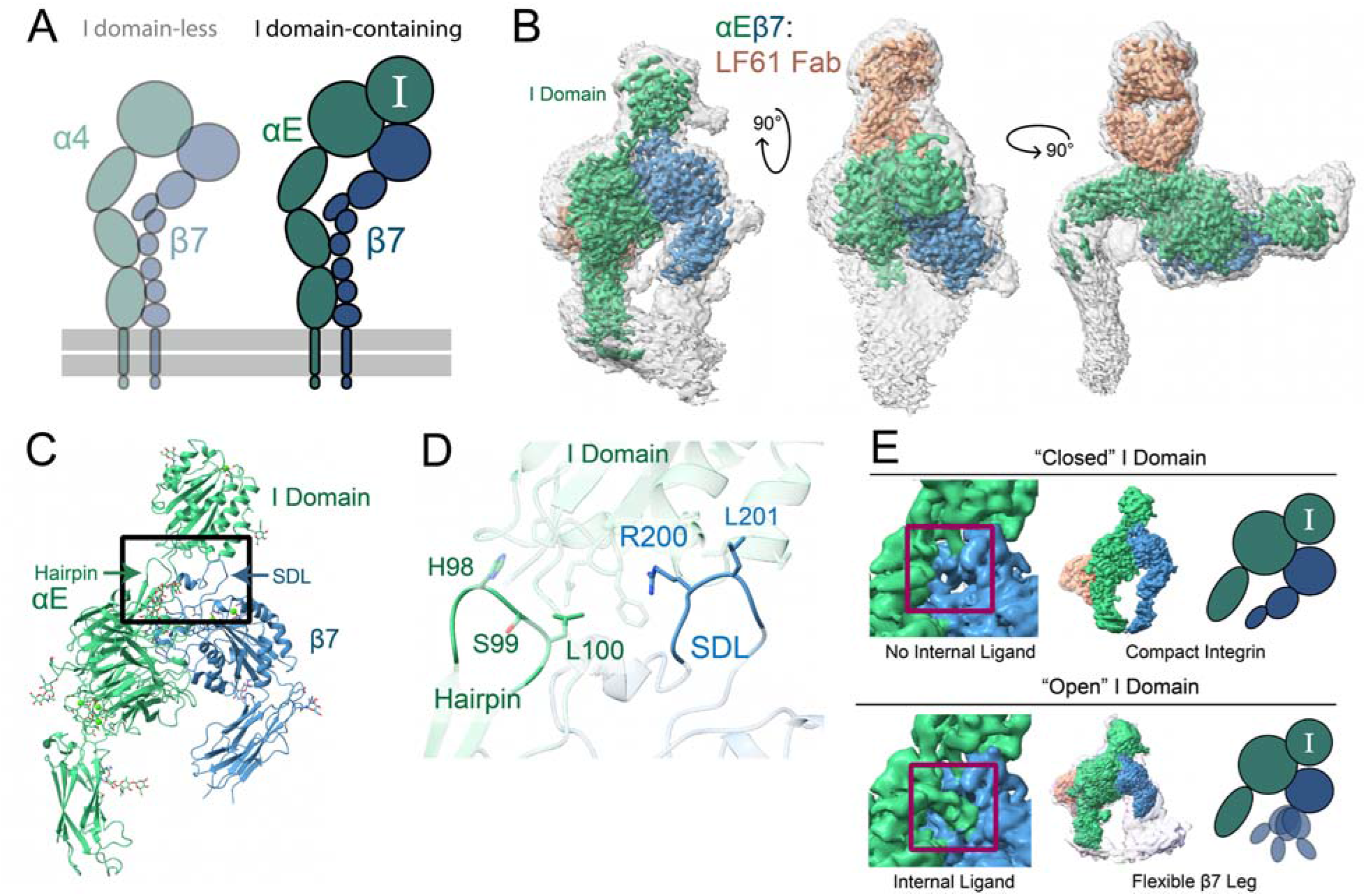
The half-bent αEβ7 inactive structure is dynamic. **A,** A schematic highlighting showing the focus of this figure on the I domain-containing integrin αEβ7. **B,** Three different views of the unsharpened αEβ7:LF61 Fab complex map. αE (green) β7 (blue) and LF61 (orange). The integrin adopts a half-bent conformation in the inactive state, and the I domain rests atop the α/β subunit interface. The colored, sharpened, high-threshold map is shown inset in the unsharpened, grayscale, low-threshold map. **C,** The refined coordinate model of the compact αEβ7 headpiece. We model five N-linked glycosylations on αE in addition to two previously reported for β7. We do not model LF61 due to a lack of sequence availability. All ions are modeled as Ca^2+^. Black box indicates regions of focus in (D-E). **D,** The αE I domain is stabilized by supports from a beta-hairpin on the αE propeller domain as well as the β7 specificity-determining loop (SDL). The αE I domain has a 940Å^2^ contact area with the rest of the integrin heterodimer, whereas the αM I domain from PDB:7P2D^51^ has only a 493.4Å^2^ contact area with the rest of αMβ2. **E,** αEβ7 assumed two broad states. In the closed I domain state (left) the C-terminal region of the α7 helix, termed the internal ligand, does not engage with β7 subunit and the β7 leg is closed inward. In the open I domain state (right) the internal ligand engages β7, leading to conformational heterogeneity in the β7 leg. For both structures the unsharpened map from a non-uniform refinement is shown in color; for the open I domain representative structures from 3D classification are shown overlaid with low opacity and a schematic is shown to the right. Red boxes indicate internal ligand region.

The I domain contains the internal ligand^12^ proposed to coordinate the ion site (βMIDAS) in its accompanying β subunit. Although our apo-αEβ7 sample was frozen in conditions that should have favored an inactive conformation, we found distinct populations of particles in the overall compact structure; some had the internal ligand engaged, whereas others did not (Fig. 3E, Supplemental Fig. 6, Supplemental Video 2). Consistent with previous analyses of isolated I domains^11^, we found that extension of the I domain’s C-terminal α7 helix is a prerequisite for internal ligand binding and βMIDAS coordination. This helix in our structure of the internally-liganded αEβ7 occupied the “open” conformation previously described in crystal structures of other I domains^33^. However, in contrast to the metastable state occupied by αXβ2^11^, the β7 hybrid domain does not prefer a single conformation upon internal ligand binding and is instead highly flexible (Fig. 3E). Our structures show that the internally-liganded state is regularly sampled by αEβ7 integrin even in conditions typically expected to favor inactive conformations.

### Ligand binding stabilizes the open **α**E I domain

To understand how movement of this I domain internal ligand is connected to external ligand binding, we next determined a cryoEM structure of the αEβ7 ectodomain bound to its primary ligand E-cadherin^34^ (Fig. 4A, Supplemental Fig. 7). At a global resolution of 3.4Å, the ligand-binding interface was well-resolved and we modeled the integrin headpiece region as well as E-cadherin EC1 (Fig. 4B). This resolves the previous uncertainty about which E-cadherin domains are necessary for integrin binding^35^. We find E-cadherin EC1 exclusively binds to the top surface of the αE I domain, using a short E-cadherin construct (EC12)^36^ (Fig. 4A). E-cadherin is known to occupy monomeric and dimeric states^37^, but we only observed it in a monomeric state when bound to αEβ7 (Fig. 4C). The αEβ7:E-cadherin molecular interface is primarily mediated by a central hydrophobic “lock and key” pocket surrounded by a network of electrostatic bonds^38^ (Fig. 4D-E). Consistent with a stable interaction, we measure a K_D_ of 51 ± 7.9 nM for the αEβ7:E-cadherin complex using Bio-Layer Interferometry (Fig. 4F). Additionally, the N303 glycosylation on the αE I domain appears to make direct contact with E-cadherin (Fig. 4G).

**Figure 4.**
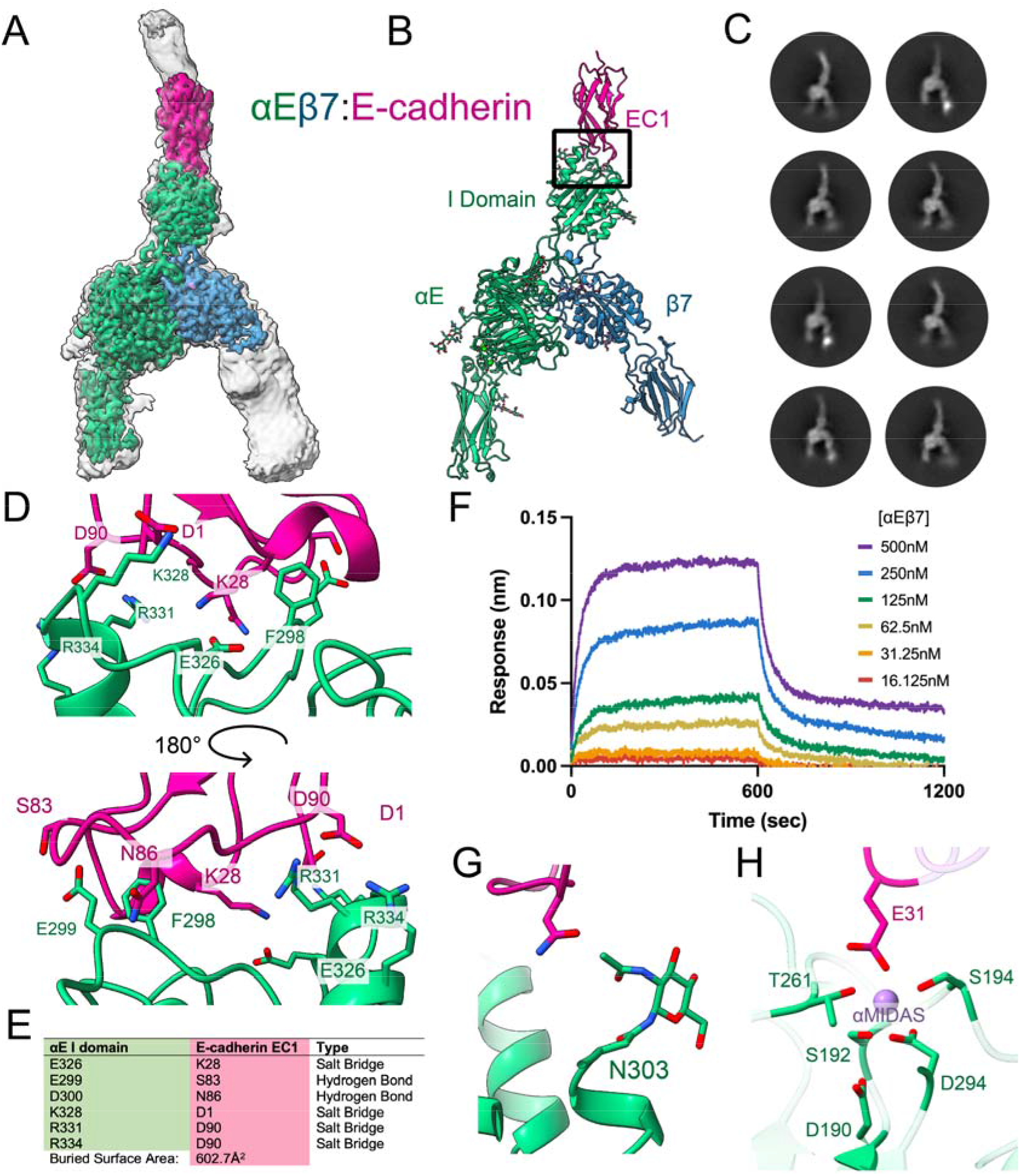
The molecular interface between integrin αEβ7 and E-cadherin. **A,** CryoEM structure of the integrin αEβ7 ectodomain (green/blue) bound to the first two EC domains of ligand E-cadherin (pink). The low threshold, unsharpened global refinement map is shown in gray, and the inset high threshold, sharpened local refinement map is shown in color. The open integrin legs suggest an active conformation. **B,** The atomic model of the αEβ7:E-cadherin complex. Upper black box indicates region of focus in (D). **C,** Representative 2D class averages of αEβ7 bound to EC12 suggest αEβ7 binds exclusively to E-cadherin monomers. **D,** The interface between αEβ7 and E-cadherin is composed primarily of a central hydrophobic residue, F298, that is surrounded by electrostatic interactions. Views are 180° rotations. **E,** Electrostatic interactions in the αEβ7:EC12 complex model between the αE I domain and EC1. **F,** Bio-Layer Interferometry (BLI) data suggest E-cadherin and αEβ7 form a high-affinity, stable interaction. **G,** The N321 glycosylation on the αE I domain contributes to ligand binding. **H,** The αE I domain (green) αMIDAS’ ion (purple) coordination sphere is completed by E-cadherin (pink) E31.

Integrin I domains contain a cation binding site (αMIDAS) coordinated by a conserved DXSXS motif. The αE I domain αMIDAS is ion-occupied in our structure, and the ion coordination complex is completed by E-cadherin E31 (Fig. 4H). As seen in crystal structures for other integrin I domains^39-41^, this αMIDAS coordination leads to an open αE I domain, with the C-terminal helix of the I domain assuming an extended conformation (Fig. 5A). Despite this large shift, the angle of the I domain relative to the apo structure remains almost entirely unchanged upon ligand binding (Supplemental Fig. 8), consistent with evidence from the compact structure that the αE I domain is particularly stable compared to other well-studied integrins. The ligand-bound αEβ7 integrin assumes an open headpiece conformation, although 3D flexibility analysis of the dataset suggests that the β7 leg maintains a large degree of free movement (Supplemental Movie 3). This movement could be crucial for integrin’s immunological function given the forces that T cells face in processes like homing, extravasation, and tissue residence^42^. Intrinsic flexibility would allow integrin to stay ligand-bound while permitting dynamic cell movement and membrane morphology changes.

**Figure 5.**
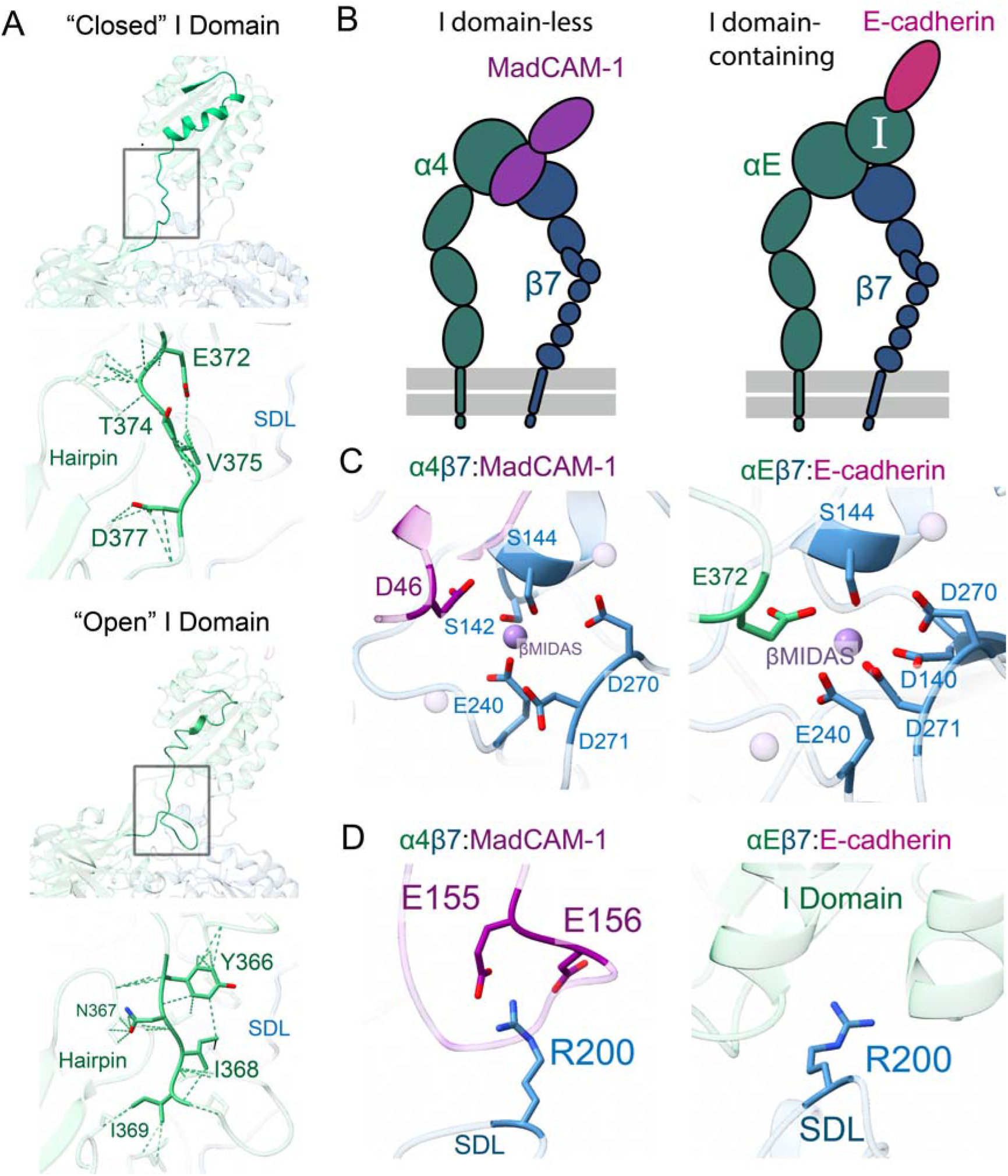
Molecular ligand mimicry by the αE I domain. **A,** The α7 helix (shown in high opacity) on the αE I domain extensively deforms upon ligand binding to allow the internal ligand to engage with β7. The closed I domain is shown top left, and the open I domain is shown bottom left. Extensive low-strength contacts stabilize the I domain’s internal ligand in either the apo closed state (top) or ligand-bound open state (bottom). “SDL” is β7 specificity-determining loop. **B,** A schematic showing the ligand-bound, activated I domain-less integrin α4β7 (left) or I domain-containing integrin αEβ7 (right). **C,** The β7 βMIDAS coordination sphere is completed in ligand bound integrins, either directly when bound to MadCAM-1 (left) or indirectly through E-cadherin-mediated allostery in αE (right). **D,** The β7 R200 residue either provides initial and stabilizing support for MadCAM-1 (left) or stabilizes the αE I domain (right).

In contrast to the unliganded structure, the I domain internal ligand is clearly engaged with β7 when bound to E-cadherin. In this open conformation, the I domain interactions shift dramatically. For example, instead of contacting residues E372-D377 of the αE propeller hairpin in the closed structure, the “collar” contacts I domain residues Y366-I369 (Fig. 5A). This shift allows the αE I domain to maintain its remarkable rigidity in both inactive and active states. This rigidity is consistent with αEβ7’s well-described role in establishing and maintaining a non-circulating, stationary tissue resident epithelial T cell population^43^. The αE E372 residue coordinates the Mn^2+^ ion occupying the β7 βMIDAS, leading to the canonical piston-like mechanism of integrin leg opening (Fig. 5C). We find these extensive collar contacts serve the biological function of stabilizing the hydrogen bonds that support the coordinating E372, an important structural feature only previously ascribed to artificial lattice contacts in prior crystal structures of open integrin I domains^11,33,41^. In our structure, this stabilization is further reinforced by the αMIDAS-mediated conformational shift in the I domain upon ligand binding as well as the presence of Mn^2+^ in the βMIDAS.

Our structures provide an unprecedented opportunity to visualize and compare conformational changes brought upon β7 either upon binding of an external ligand in the α4β7 non-I domain configuration, or by an internal ligand in an I-domain containing αEβ7 heterodimer (Fig. 5B). The same βMIDAS ion is coordinated by αE E372 in the αEβ7 heterodimer as by MadCAM-1 D46 within the α4β7:MadCAM-1 complex (Fig. 5C). This leads to the same open conformation and leg flexibility in α4β7 as seen in internally-liganded αEβ7. The RMSD of the β7 βI domain in ligand-bound α4β7 vs. αEβ7 is 0.6Å, indicating that β7 undergoes nearly identical behavior in the context of either α4 or αE when activated. Remarkably the β7 SDL also uses the exact same arginine, R200, to support MadCAM-1 as it does the αE I domain (Fig. 5D). Collectively these data provide unambiguous support for the structural mimicry of external ligand-mediated integrin activation by the intrinsic I domain at key interaction sites.

### Collagen origin of the integrin I domain

The evolutionarily ancient I domain insertion not only expanded the ligand-binding repertoire of integrin proteins but did so without perturbing the ancestral intricate conformational changes required for activation. We wondered if the I domain’s ancestor already contained all the necessary machinery to facilitate the allosteric communication seen in our structures. Sequence alignments show the I domain was inserted immediately following a structural proline between beta-propeller blades 2 and 3 (Supplemental Fig. 1B). To find the common ancestor of all I domains, we aligned a representative subset of integrin I domains from across the *Olfactores* phylogeny. We used this alignment to generate a Hidden Markov Model (HMM) of the protein domain encompassing the I domain and internal ligand (Supplemental Fig. 9). Finally, we searched the genomes of cephalochordates, the direct outgroup of *Olfactores*, with this I domain HMM. This strategy enabled us to identify homology to the presumed I domain ancestor rather than the multiple other vWFA domains present in animal genomes. Our analyses reveal that the most likely ancestors of the α-integrin vWFA-like I domains are vWFA domains from extracellular matrix proteins (Fig. 6A). More specifically, we observe that vertebrate integrin I domains phylogenetically cluster within the alpha-3(VI) class of collagens from cephalochordates to the exclusion of all other vWFA domains (Fig. 6B).

**Figure 6.**
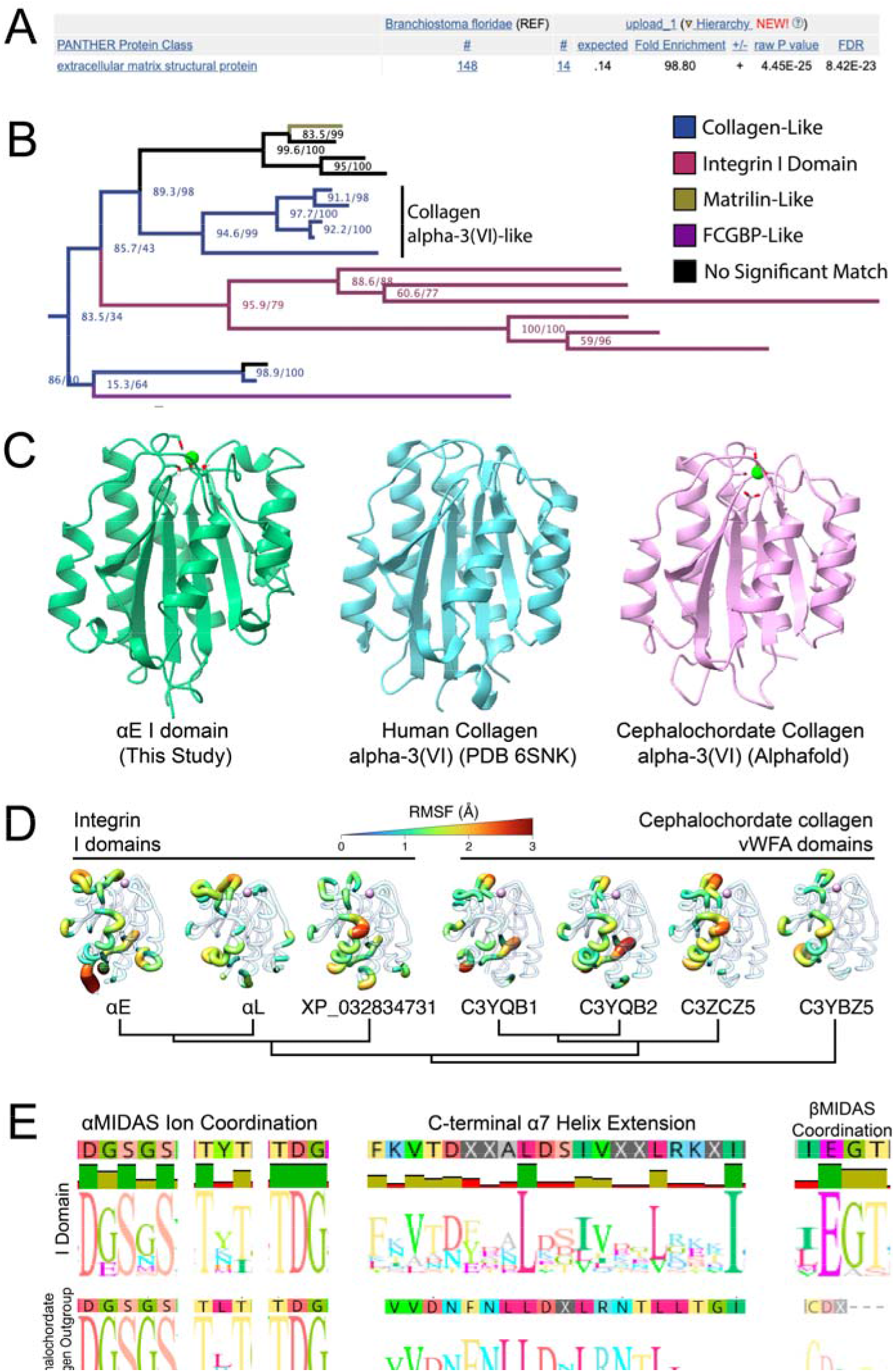
The integrin I domain was co-opted from an ancient collagen gene. **A,** A PANTHER protein classification analysis of the top *Branchiostoma floridae* hits from our HMM search shows that extracellular matrix proteins are significantly over-represented. **B,** A phylogenetic tree of the VWF-like domains of the top cephalochordate hits from our HMM search was generated using IQ-TREE. The full cephalochordate protein sequences from each hit were reciprocally BLASTed against the human genome to identify protein homology, and those results are displayed in the colors shown. The cutoff for “no significant match” is an E value < 1e-30. Node values are displayed as “ultrafast bootstrap/SH-aLRT”. The integrin I domain is most closely related to VWF-like domains from cephalochordate collagen alpha-3(VI)-like proteins. **C,** Atomic models of the closed I domain (this study), a human collagen alpha-3(VI) (PDB 6SNK^43^) VWF-like domain, and an alphafold prediction of a VWF-like domain of *Branchiostoma floridae* collagen alpha-3(VI) (Uniprot C3YQB2) show high visual structural similarities. **D,** Molecular dynamics (MD) simulations of representative collagen or integrin vWFA domains bound with Mg^2+^ in the MIDAS site suggest characteristic dynamics were present in the common integrin I domain ancestor. The individual residue flexibility (i.e. root mean square fluctuations, RMSF) is projected on the three-dimensional structure and colored from 0 (blue) to >3 (red). Regions of fluctuations greater than 1.0 Å are highlighted. **E,** Logo plots of alignments of either representative I domains (top) or homologous domains from outgroup cephalochordate collagens (bottom) show that many important sequence features were likely present in the integrin I domain common ancestor. The βMIDAS ion-coordinating glutamate (right) was not found in outgroup sequences.

Based on our phylogenetic analyses, we conclude that the original I domain acquisition in α-integrins was derived from a collagen domain involved in collagen-collagen interactions. Outgroup cephalochordate proteins and human collagen VI both share high structural homology with the αE I domain (Fig. 6C). Microseconds-long molecular dynamics simulations reveal that key dynamic movements present in the I domain were also present in this collagen ancestor (Fig 6D, Supplemental Fig 10). Collagen proteins commonly form oligomeric structures within or between collagen types, including collagen alpha-3(VI)^44^. Based on our analyses, we infer that the ancestral I domain was capable of binding collagen immediately upon acquisition, providing many functionally relevant motifs for integrin I domain function, including those involved in ion coordination and in structural changes in the C-terminal helix (Fig. 6E). The MIDAS performs critical functions in collagens which contain it^45,46^, just as it does in integrin I domains.

However, this acquisition could not explain the presence of the highly conserved I/LEGT motif containing the βMIDAS-coordinating glutamate required for internal ligand-based activation (Fig. 6E). Indeed, we found no significant matches to this motif either within collagen or within any other vWFA proteins in ProtKB metazoan genomes. This suggests either of two possibilities. The first possibility is that the I/LEGT motif was a distinct acquisition from an independent evolutionary ancestor that coincided with the insertion of the I domain into the ancestral integrin gene. However, the probability of precisely acquiring the exact sequence needed to render the I domain functional from a separate source seems highly unlikely. Alternatively, the motif might have been already present in an ancestral α-integrin gene but was not under stringent selective constraints until after the acquisition of the I domain. In support of this possibility, multiple extant α-integrins lacking I domains nonetheless encode EG motifs within the flexible loop between the second and third beta propeller blades (Supplemental Fig 11A). Although these motifs are not conserved between subunits (Supplemental Fig 11B), we posit that the presence of these otherwise unconserved, prototypical EG motifs became selectively entrenched in the α-integrin that acquired the I-domain insertion from collagen, once the function of the I/LEGT motif became critical for integrin activation.

### Concluding remarks

Here we use cryoEM to describe the strategy by which the intrinsic integrin I domain structurally mimics an extrinsic integrin ligand. We leverage the β7 integrins, the I domain-containing αEβ7 and I domain-lacking α4β7, to determine structures of both integrins alone and engaged with their ligands: E-cadherin and MadCAM-1, respectively. Using these structures, we map out these ligand-binding interfaces and define the key structural features involved in this I domain-based activation mechanism. Integrins have characteristic conformational heterogeneity that historically has made them difficult to study structurally at high resolution; we have used recent computational advances to resolve sample heterogeneity and determine structural ensembles, which has allowed us to characterize the complicated allosteric relay facilitated by this evolutionary insertion. Furthermore, by defining the molecular features of the β7 integrin family we provide the necessary foundation for generating targeted, rationally-designed therapeutics that will be highly applicable to a range of diseases from the IBD family to cancer.

Our results detail a structural framework for how a high-risk, high-reward evolutionary insertion can preserve necessary protein functions while serving as a launch pad for adaptation. Despite completely occluding integrin’s ancestral ligand binding site, the integrin I domain remarkably retains the same functional output of an ancestral integrin ligand at multiple levels. In this way, the I domain served two capacities at its inception: the ability to bind to an ion-coordinating ligand, as well as to coordinate downstream integrin conformational changes. Given the collagen origin of the I domain itself, it is likely that the I domain’s co-option into the integrin family catalyzed the ability of integrins to directly associate with the collagen extracellular matrix (Fig 7). This ancestral feature^47,48^ has greatly expanded and specialized in extant integrins^49^. Given that most eukaryotic proteins contain multiple distinct structural domains^50^, we anticipate this form of evolutionary molecular co-option provided novelty across a wide array of molecular processes. Collectively, our analyses describe the unique and shared structural features of I domain-mediated integrin activation that were critical for its evolutionary and immunological success.

**Figure 7.**
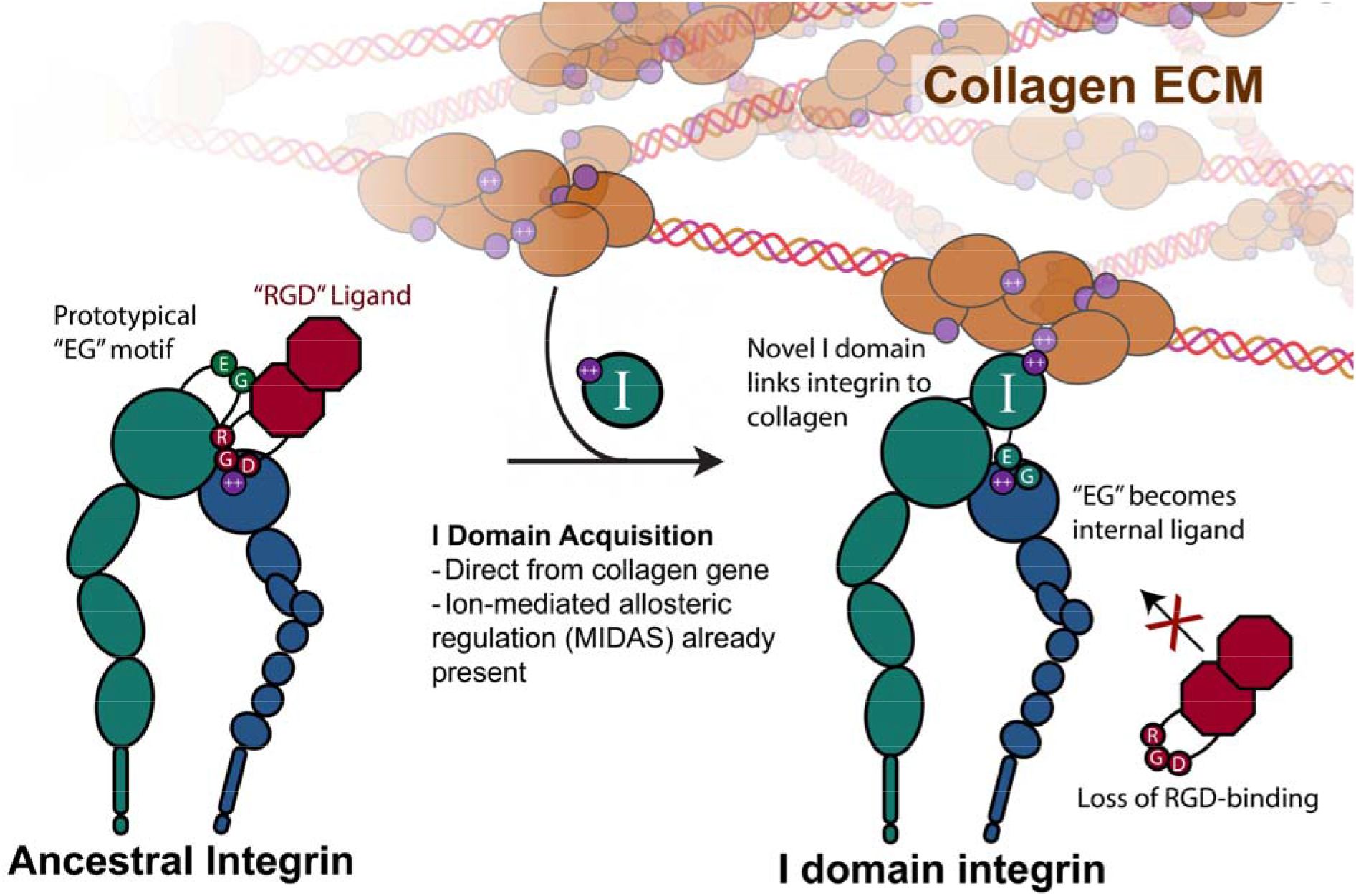
A model for the evolutionary origin of the novel integrin I domain. The ancestral state integrin binds ion-coordinating ligands directly at the heterodimer interface on the basis of a conserved motif. The derived vWFA-family integrin I domain was acquired directly from an ancient collagen gene, which immediately endowed the new I domain-containing integrin with the ability to directly interact with the collagen extracellular matrix, at the cost of losing the specific sequence recognition. We propose the necessary parts for this acquisition to be successful were all present prior to its co-option, including a sequence that would become the ion-coordinating activation mechanism (the internal ligand or ligand “mimetic”), as well as the ion-mediated allosteric relay within the I domain itself.

## Methods

### DNA Constructs

The region encoding the wiltype αE ectodomain (M1-H1123, used for integrin:e-cadherin complex) followed by a C-terminal linker, HRV 3C cut site (LEVLFQGP), acidic coil motif (AQCEKELQALEKENAQLEWELQALEKELAQ) and Strep-Tag II (WSHPQFEK*) inserted into a pcDNA3.1-Hygro(-)-like backbone was synthesized commercially (GenScript). The R177G/R178G, “RRtoGG,” mutant (clasped construct used for integrin:Fab complex) was generated via site-directed mutagenesis (NEB). The region encoding the β7 ectodomain (M1-H723) followed by a C-terminal linker (GTGG), HRV 3C cut site (LEVLFQGP), basic coil motif (AQCKKKLQALKKKNAQLKWKLQALKKKLAQ) and 6xHIS tag inserted into a pcDNA3.1-Hygro(-)-like backbone was synthesized commercially (GenScript). The region encoding the α4 ectodomain (M1-Q970) was amplified from Addgene plasmid #81178 and inserted into a pcDNA3.1-Hygro(-)-like backbone using standard molecular cloning techniques. The region encoding the MadCAM-1 ectodomain (M1-Q317) was synthesized commercially as a G-block (IDT) and inserted into a pcDNA3.1-Hygro(-) with the same C-terminal linker, 3C cut site, basic coil motif, and 6xHIS tag as β7 using standard cloning techniques. For Fc-tagged E-Cadherin, the region encoding the E-cadherin ectodomain (D155-A698) with a CD33 signal sequence (GMPLLLLLPLLWAGALA) (Gift from Barry Gumbiner) was inserted into a pcDNA3.1-Hygro(-)-like with a C-terminal HRV 3C cut site and human Fc tag amplified from Addgene plasmid 145164 using standard cloning techniques. The bacterial expression plasmid encoding an N-terminal 6xHIS-SUMO-tagged EC12 protein (E-cadherin EC domains 1 and 2, D155-N371) was a gift from Barry Gumbiner.

### Protein expression and purification

Both integrins αEβ7 and α4β7, as well as MadCAM-1, were expressed in mammalian ExpiCHO cells (ThermoFisher). The basic expression and purification protocol is similar for each protein. Protein expression was performed according to the “Max Titer’’ manufacturer recommendations. For each integrin, a ratio of 3:2 α:β DNA was transfected for a total of 1µg of DNA per mL of cell culture. For MadCAM-1 a total of 1µg of DNA per mL of cell culture was used for transfection. Cells were grown to a concentration of 7-10 x 10^6^ cells/mL at 37°C, 8% CO_2_, and 90% humidity. The day of transfection, cells were split to 6 x 10^6^ cells/mL and allowed to recover for 2-4 hours. Cells were then transfected according to manufacturer recommendations using Expifectamine CHO and grown overnight at 37°C, 8% CO_2_, and 90% humidity. The next day, cells were given Enhancer and Feed according to manufacturer recommendations and shifted to 32°C, 5% CO_2_, and 90% humidity to support protein expression. Five days post-transfection, cells were given a second dose of Feed according to manufacturer recommendations. 8-10 days post-transfection, cells were harvested via centrifugation for 15 minutes at 4°C and 1000xg. The cell supernatant containing secreted protein was further clarified for 20 minutes at 4°C and 30000xg and diluted 2:1 into HisTrap Binding Buffer (20mM NaPO_4_, 500mM NaCl, 20mM imidazole, pH 7.4). The clarified supernatant was flowed over a 5mL HisTrap FF Crude column (Cytiva) equilibrated in HisTrap Binding Buffer using an ÄKTA Pure 25 L1 FPLC system. Bound protein was eluted in HisTrap Elution Buffer (20mM NaPO_4_, 500mM NaCl, 500mM imidazole, pH 7.4). Fractions containing protein according to A_280_ were pooled, concentrated to 500µL, and buffer-exchanged into integrin storage buffer (20mM Tris-HCl pH 7.4, 150mM NaCl, 1mM MgCl_2_, 1mM CaCl_2_) using an Amicon Ultra-15 concentrator (Millipore) with a 10kDa cutoff. For clasped integrin proteins, the concentrate was then purified via gel filtration chromatography using a Superdex 200 Increase 10/300 SEC column (Cytiva) equilibrated with integrin storage buffer. Peak fractions containing integrin according to SDS-PAGE were pooled, concentrated to 2mg/mL, snap frozen in 10% (v/v) glycerol and stored at - 80°C. For MadCAM-1 and unclasped integrins, 1:10 (w/w) of 3C protease was added to the protein concentrate and incubated overnight with end-over-end rotation at 4°C. The next day the cleaved protein was further purified via gel filtration chromatography, concentrated, and stored as was done with the clasped protein.

Human Fc-tagged E-cadherin ectodomain was expressed and purified similarly to integrin and MadCAM-1 constructs. Ectodomains were expressed in ExpiCHO cells according to the “Max Titer’’ manufacturer recommendations using a total of 1µg of DNA per mL of cell culture. Cells were fed and harvested and supernatants were clarified as with the integrin constructs. Supernatant was diluted 2:1 in HiTrap binding buffer (50mM Tris-HCl pH 7.4, 150mM NaCl, 3mM CaCl_2_) and flowed over a 1mL HiTrap Protein G HP column (Cytiva) equilibrated in HiTrap binding buffer. The column was washed with 10 volumes of HiTrap binding buffer, and protein was eluted with 100mM glycine-HCl, pH 2.7 into 1M Tris-HCl, pH 9.0 to neutralize. Eluate was pooled and concentrated to 500µL, and was then further purified via size exclusion chromatography on a Superdex 200 Increase 10/300 SEC column (Cytiva) equilibrated in 50mM Tris-HCl pH 7.4, 150mM NaCl, 3mM CaCl_2_. Peak fractions were pooled, concentrated to 2mg/mL, snap frozen with 10% (v/v) glycerol and stored at -80°C.

EC12 was expressed and purified as previously described^58^ with slight modifications. Briefly, BL21 DE3 cells containing the expression plasmid were grown to an OD_600_ of 0.6, induced with 0.1mM IPTG, and incubated with shaking overnight at 18°C. The following day cells were harvested at 4000xg for 30 minutes at 4°C and resuspended in 10mL/L of cell culture in EC12 binding buffer (500mM NaCl, 20mM Tris-HCl pH 7.4, 3mM CaCl_2_, 20mM imidazole) with EDTA-free cOmplete protease inhibitor cocktail (Roche). The cells were lysed via sonication on ice for 3 minutes and the lysate was clarified for 30 minutes at 4°C, 20000xg. The lysate was flowed over a 1mL HisTrap FF Crude column (Cytiva) equilibrated in EC12 Binding Buffer and washed with 10 CV of EC12 Binding Buffer. Bound protein was eluted in 500mM NaCl, 20mM Tris-HCl pH 7.4, 3mM CaCl_2_, 250mM imidazole. Peak fractions were pooled and concentrated using an Amicon Ultra-15 concentrator (Millipore) with a 10kDa cutoff. The concentrate was dialyzed overnight at 4°C into 4L of 50mM Tris-HCl pH 7.4, 150mM NaCl, 3mM CaCl_2_ with 250 units of 6xHIS-tagged SUMO protease (ThermoFisher) for scar-free SUMO tag cleavage. The following day the dialysate was run over a 1mL HisTrap FF Crude column to remove the protease and unprocessed EC12. The dialysate was then further purified via gel filtration chromatography on a Superdex 75 Increase 10/300 GL (Cytiva). Peak fractions containing EC12 were pooled, concentrated to 2mg/mL, and snap frozen in 10% v/v glycerol. We attempted to express C-terminally HIS-tagged EC12 in mammalian cell culture but proteins consistently precipitated following HisTrap elution.

### Negative-stain EM sample preparation

Integrin αEβ7 was diluted to a final concentration of 2-20µg/mL in buffer containing 20mM Tris-HCl pH 7.4, 150mM NaCl, and ion concentrations indicated in Supplemental Fig 5. 3µL of diluted integrin was applied to a glow-discharged 400 mesh copper glider grid (Ted Pella) that had been covered with a thin layer of continuous amorphous carbon. The grids were stained with a solution containing 2% (w/v) uranyl formate as previously described^59^.

### Negative-stain EM data acquisition and processing

Data were acquired using a Thermo Fisher Scientific Talos L120C transmission electron microscope operating at 120 kV and recorded on a 4k x 4k Thermo Scientific Ceta camera at a nominal magnification of 92000x with a pixel size of 0.158 nm. Leginon^60^ was used to collect 596 (5mM CaCl_2_), 187 (1mM MgCl_2_, 1mM CaCl2), or 262 (1mM MnCl_2_) micrographs at a nominal range of 1.5-2.5 µm under focus and a dose of approximately 50 e^-^/Å^2^.

Similar processing pipelines were used for all negative stain datasets. Micrographs were processed using GCTF^61^, Gautomatch (https://github.com/JackZhang-Lab) and cryoSPARC^62^. CTF estimation of micrographs for the 5mM CaCl_2_ and 1mM MgCl_2_, 1mM CaCl_2_ conditions was performed using GCTF. Initially, 93294 (5mM CaCl_2_) or 73949 (1mM MgCl_2_, 1mM CaCl_2_) particles were picked in a reference-free manner using Gautomatch and imported into cryoSPARC. The particles were subjected to 2 rounds of 2D classification for a final particle count of 17199 (5mM CaCl_2_) or 12289 (1mM MgCl_2_, 1mM CaCl_2_) contributing to 2D classes. CTF estimation of micrographs for the 1mM MnCl2 condition was performed using Patch CTF in cryoSPARC. An initial stack of 109907 particles picked using the blob picker in cryoSPARC was subjected to 2 rounds of 2D classification for a final stack of 36469 particles contributing to 2D averages.

### cryoEM sample preparation

The apo α4β7 protein was diluted 1:10 in non-activating buffer (20mM Tris-HCl pH 7.4, 150mM NaCl, 5mM CaCl_2_) to a final concentration of 0.2mg/mL. The RRtoGG αEβ7 ectodomain was diluted 1:10 in non-activating buffer and incubated with digested LF61 Fab (Invitrogen) at a 1:4 molar ratio for 45 minutes at room temperature with end-over-end mixing. The integrin:Fab complex was subject to size exclusion chromatography in non-activating buffer and complex peaks were concentrated to 0.25 mg/mL. The wildtype αEβ7 ectodomain was diluted 1:10 in activating buffer (20mM Tris-HCl pH 7.4, 150mM NaCl, 1mM MnCl_2_) and incubated with EC12 at a 1:4 molar ratio for 45 minutes at room temperature with end-over-end mixing. The integrin:ligand complex was subject to size exclusion chromatography in activating buffer and complex peaks were concentrated to 1 mg/mL. Immediately before freezing, stock CHAPS detergent was added to the integrin:ligand complex to a final concentration of 0.05% (v/v). The α4β7:MadCAM-1 complex was formed and prepared in the same way as αEβ7:EC12. Following dilution and complex formation steps, all samples were frozen with similar conditions. 3µL of protein was applied to Quantifoil grids (EMS) that were glow-discharged for 30s at 15 mA. The grids were blotted with a Vitrobot Mark IV (ThermoFisher) using a blot time of 3-7s and blot force of 4-5 at 100% humidity and 4°C. Grids were plunge-frozen in liquid ethane cooled by liquid nitrogen and stored in liquid nitrogen.

### CryoEM data acquisition and processing

The details of datasets and processing pipelines are outlined in Table 1, Supplemental Figure 2 (α4β7:MadCAM-1), Supplemental Figure 4 (apo α4β7), Supplemental Figure 6 (αEβ7:LF61 Fab) and Supplemental Figure 7 (αEβ7:EC12). All datasets were collected on a Glacios cryo-transmission electron microscope (ThermoFisher) operating at 200 kV and recorded with a Gatan K3 Direct Detection Camera. Automated data collection was carried out using SerialEM^63^. One hundred-frame movies were recorded in super-resolution counting mode at a nominal magnification of 36000x corresponding to a calibrated super-resolution pixel size of 0.561 Å/px. Each dataset was collected with a nominal defocus range of 1.2-1.8 µm under focus and a dose of approximately 50 e^-^/A2.

Dose fractionated super-resolution movies were motion-corrected and binned 2x2 by Fourier cropping using MotionCor2^64^ within the RELION^65^ wrapper. From there, motion-corrected stacks were further processed in cryoSPARC according to each processing pipeline. Briefly, particles were picked using the blob picker, subjected to multiple rounds of 2D and/or 3D classification using *ab initio* and heterogeneous refinement using initial models generated from the data, and further refined with non-uniform and local refinement. Masks for local refinements were generated using UCSF ChimeraX^66^ and cryoSPARC. Map sharpening was performed using the Cosmic2^67^ server wrapper of DeepEMhancer^68^ with the high resolution setting. Local resolution estimation was performed in cryoSPARC. 3D FSC^69^ plots were generated from https://3dfsc.salk.edu/.

CryoSPARC was used for all motion and variability analyses. We used the cluster setting in 3D Variability Analysis^15^ followed by homogeneous and heterogeneous refinement to sort the αEβ7:LF61 Fab particles into open or closed I domain states. Maps showing the conformational range of the internal-liganded β7 leg in the Fab-bound structure (Fig 1D) were generated using 3D Classification in PCA mode. We generated the volume series showing the continuous motion of the β7 in ligand-bound structures using 3DFlex^14^. All map visualizations (images and movies) were generated and recorded using UCSF ChimeraX.

### Model Building

For the Fab-bound initial αEβ7 model we used an Alphafold2^70^ model of αE and PDB: 3V4P^24^, Chain B for β7. We do not model the LF61 Fab due to a lack of sequence availability. The initial models were manually fit into the density using UCSF ChimeraX, followed by dock-in-map in Phenix^71^. For the initial αEβ7:EC12 model we used the alphafold αE model, a homology model of β7 built in SWISS-MODEL^72^ using PDB: 7NWL^31^, chain B as the reference, and PDB: 4ZT1^73^, Chain A for EC12. We generated the initial α4β7:MadCAM-1 model using alphafold-multimer^74^.

Models were manually adjusted in an iterative way between COOT^75^ and ISOLDE^76^ within UCSF ChimeraX. The β7 hybrid domain was flexibly fit into the ligand-bound models using ISOLDE. The αEβ7:LF61 Fab model was refined using the closed I domain map. Glycans were built manually in COOT using the “Carbohydrate” module. Models were built using a combination of the non-uniform and locally-refined, unsharpened and sharpened maps. Rosetta^77^ was used to estimate B-factors and depict secondary structure. All maps used for model building have been deposited.

Electrostatic molecular interactions were determined using the PISA server^57^. RMSD calculations for the βI domain were done in ChimeraX using the “matchmaker” function with β7 residues V133-S369.

### Molecular Dynamics Simulations

To understand the ancestral conformational dynamics of the integrin I domain, we performed all-atom molecular dynamics simulations on a representative sample of integrin I domains (αE, αL, α2, *Petromyzon marinus* XP_032834731), sister clade cephalochordate collagen-derived vWFA domains (C3YQB1, C3YQB2, C3ZCZ5, C3YBZ5, C3Y2U7), and RPN10 to be used as a distant outgroup. Simulations were performed using OpenMM 7.7.0^78^ employing AMBER19ffSB force field^79^, OPC water model, and 12-6-4 Li Merz ion parameters^80^. The initial coordinates of the studied I domains were obtained from the cryo-EM model (αE: this study), prior crystal structures (αL: 3F74^81^, α2: 5HJ2^82^, RPN10: 5LN1^83^), or AlphaFold2 predicted models (C3YQB1, C3YQB2, C3ZCZ5, XP_032834731, C3YBZ5, C3Y2U7). AlphaFold2 predictive models were generated in Colabfold^84^.

The proteins were solvated in a periodic truncated octahedron with a minimum distance of 16Å on all sides and 150mM NaCl. One magnesium, manganese, or calcium ion was placed in the MIDAS site. For RPN10, no additional ion was modeled. Prior to production, systems were energy minimized for 20,000 steps and equilibrated at 298K with backbone atoms harmonically restrained by a 5 kcal/mol*Å^2^ force constant for 10 ns. Langevin dynamics was performed using a Langevin integrator with an integration timestep of 4 fs and collision rate of √2 ps^-1^ and hydrogen mass repartitioning^85^. Pressure was maintained using a Monte Carlo barostat with an update frequency of 100 steps. Nonbonded interactions were calculated with a distance cutoff of 10Å. Trajectory snapshots were saved every 100 ps during production simulations. For each ion condition (Mg^2+^, Mn^2+^, Ca^2+^), ten simulation repeats with different initial velocities were conducted, each 2.4µs in length, totaling in ∼670µs of aggregate simulation time.

Trajectories were processed using in-house scripts utilizing CPPTRAJ^86^ and MDTraj^87^ packages. The last 2.0µs of each simulation trajectory was used for analysis. RMSD and RMSF measurements were performed on Cα atoms and using the initial structure for simulation as the reference.

### Bio-Layer Interferometry (BLI)

Assays were performed on an Octet Red (ForteBio) instrument at 25°C with shaking at 1,000 RPM. Protein A biosensors were hydrated in activating buffer with the addition of 0.1% BSA and 0.02% Tween-20 (BLI buffer) for 15 min. Fc-tagged E-cadherin ectodomain was loaded at 20 μg/mL in BLI buffer until a threshold of 1.2nm was reached. A baseline equilibration step was performed in BLI buffer for 2 min. Association of αEβ7 in BLI buffer at various concentrations in a two-fold dilution series from 500nM to 16.125nM was carried out for 600 s prior to dissociation for 600 s. The data were baseline subtracted prior to fitting performed using a 1:1 binding model and the ForteBio data analysis software. Mean K_D_ values were determined with a global fit applied to all data from three independent replicates. A representative sensorgram of the three replicates is displayed in the relevant figure.

### Sequence curation and Hidden Markov Model (HMM) searching

Protein sequences for all human integrin subunits were curated in the NCBI database wrapper within Geneious Prime. Representative I domain sequences from tunicates were curated using BLAST with a urochordate taxon restriction and human αL as a search query. Hidden Markov Model (HMM) searching of UniProtKB was performed on the online HMMER server from the European Bioinformatics Institute^88^ with the full integrin I domain alignment as an initial query. Since no sequences containing the internal ligand I/LEGT motif were found, we restricted our search to include only residues 1-160 of the I domain alignment and included only results from *Branchiostoma*. Sequences with an individual E-value of less than 1e-30 were considered hits and used for subsequent alignment and phylogenetic analyses.

### Sequence alignments

All protein sequence alignments were generated using the default MUSCLE algorithm^89^ and visualized using Geneious Prime. Alignments were further curated manually to remove large gaps and sequences with low quality. Sequence alignments are available as supplemental text files in FASTA or PHYLIP format with the following names: Alignment of all human β subunits (Supplemental Data 1), Alignment of all human α subunits (Supplemental Data 2) except αE which was excluded due to an extra “X” domain preceding the I domain that is unique to αE, Alignment of integrin I domains across humans and tunicates used for HMM search (Supplemental Data 3), Alignment of all hits from HMM search (Supplemental Data 4). Sequence names are provided as protein names, NCBI accession numbers or Uniprot identifiers.

### Phylogenetic Analysis

All phylogenetic trees were generated using a maximum likelihood framework within IQ-TREE^90^. Substitution models for each alignment were estimated with IQ-TREE. Both 1000 ultrafast bootstrap and SH-aLRT replicates are displayed as branch support statistics, also generated in IQ-TREE. Trees were visualized in FigTree^91^. For the tree presented in Fig 6 the topology was constrained such that the integrin I domains form a monophyletic clade.

### PANTHER Functional Analysis

Protein functional classification was performed via the online PANTHER database classification system (http://pantherdb.org/) using the significant *Branchiostoma floridae* Uniprot hits. Half of the included sequences were classified as extracellular matrix-related, while the other half were functionally unclassified.

## Acknowledgments

Funding for this research was supported by the National Institutes of Health under Grant R35 GM147414. This work was in part funded by a National Science Foundation Graduate Research Fellowship and NIH T32GM008268 (JAH) and a Pew Biomedical Scholars award (MGC). This work was made possible with the support of the Mahan Fellowship to MCC. This fellowship is made possible by funding from Mark and Nikki Mahan. Electron microscopy data were generated using the Fred Hutchinson Cancer Center Electron Microscopy shared resource, supported in part by the Cancer Center Support Grant P30 CA015704-40. We would like to thank Theo Humphreys, Dr. Anvesh Dasari, and Steve MacFarlane for their microscopy assistance and knowledge. We are grateful to Dr. Barry Stoddard, Dr. Thamiya Vasanthakumar, Rachel Werther, and Andres Fernandez for comments on the manuscript. We would like to thank Adam Nguyen, Rachel Werther and Dr. Caleigh Azumaya for invaluable assistance with training on biochemical and structural techniques. The funders played no role in the study design, data collection and interpretation, or the decision to publish this study. HSM is an Investigator of the Howard Hughes Medical Institute. This article is subject to HHMI’s Open Access to Publications policy. HHMI lab heads have previously granted a nonexclusive CC BY 4.0 license to the public and a sublicensable license to HHMI in their research articles. Pursuant to those licenses, the author-accepted manuscript of this article can be made freely available under a CC BY 4.0 license immediately upon publication.

## Author contributions

J.A.H., M.C.C., H.S.M., and M.G.C. conceived the project. J.A.H. performed biochemical purification, kinetics assays, electron microscopy data collection and processing, model building, and phylogenetic experiments. M.C.C. performed molecular dynamics simulations and analysis. H.S.M. and M.G.C. supervised the project. J.A.H., M.C.C., H.S.M., and M.G.C. acquired funding for the research. J.A.H., M.C.C., and M.G.C. wrote the original draft. J.A.H., M.C.C., H.S.M., and M.G.C. edited and finalized the manuscript.

## Data availability

All generated atomic protein models, cryoEM density maps, and motion-corrected micrographs have been deposited in the Protein Databank (PDB), Electron Microscopy Databank (EMDB), and Electron Microscopy Public Image Archive (EMPIAR) respectively. The α4β7:MadCAM-1 structure is assigned PDB-XXX, EMDB-XXXX, and EMPIAR-XXXX. The apo-α4β7 structure is assigned EMDB-XXXX and EMPIAR-XXXX. The αEβ7:LF61 closed I domain Fab structure is assigned PDB-XXXX, EMDB-XXXX, and EMPIAR-XXXX. The αEβ7:LF61 open I domain structure is assigned EMDB-XXXX and EMPIAR-XXXX. The αEβ7:EC12 structure is assigned PDB-XXXX, EMDB-XXXX, and EMPIAR-XXXX. Sequences with accompanying protein alignments for evolutionary analyses are provided as supplemental data files. All other data are available from authors upon request.

---

**Supplemental Figure 1.**
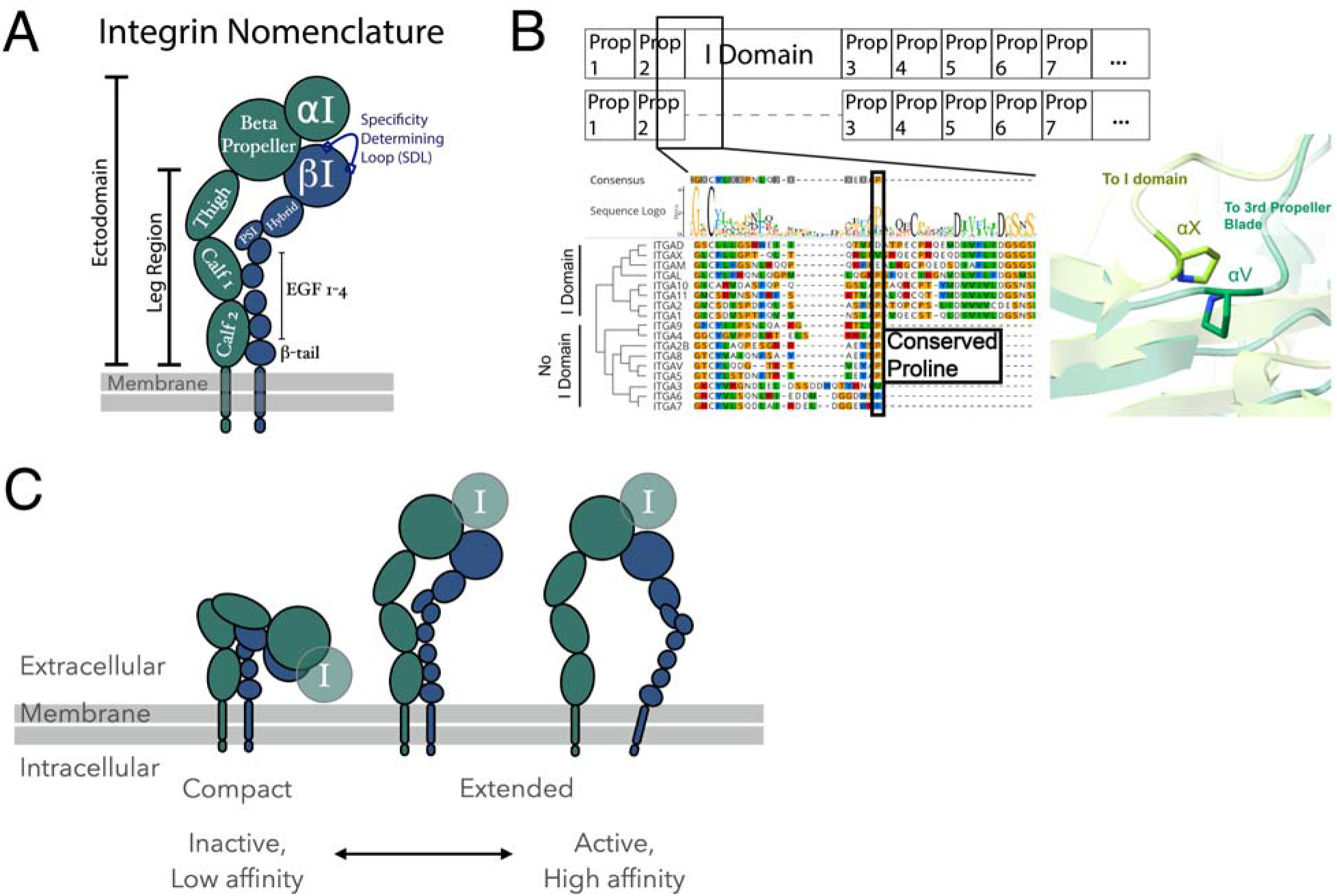
Structural features of integrin molecules. **A,** A schematic of integrin ectodomain domain and the beta-propeller organization. The I domain has inserted between the second and third of seven β-propeller domain repeats in some α-integrin genes. **B,** An alignment of all human integrin α subunits (except αE) shows insertion of the I domain in the ancestral integrin gene occurred immediately following a conserved proline at the end of the second β-propeller blade. Slightly offset structural overlays of the I domain-less integrin αV (dark green, PDB 1L5G^52^) and I domain-containing αX (lime green, PDB 4NEH^11^) with their respective conserved prolines displayed. **C,** Integrin conformations range from a compact, inactive state with low ligand affinity (left) to an extended-open, active state with high ligand affinity (right). Approximate hypothesized I domain locations are shown as semi-transparent.

**Supplemental Figure 2.**
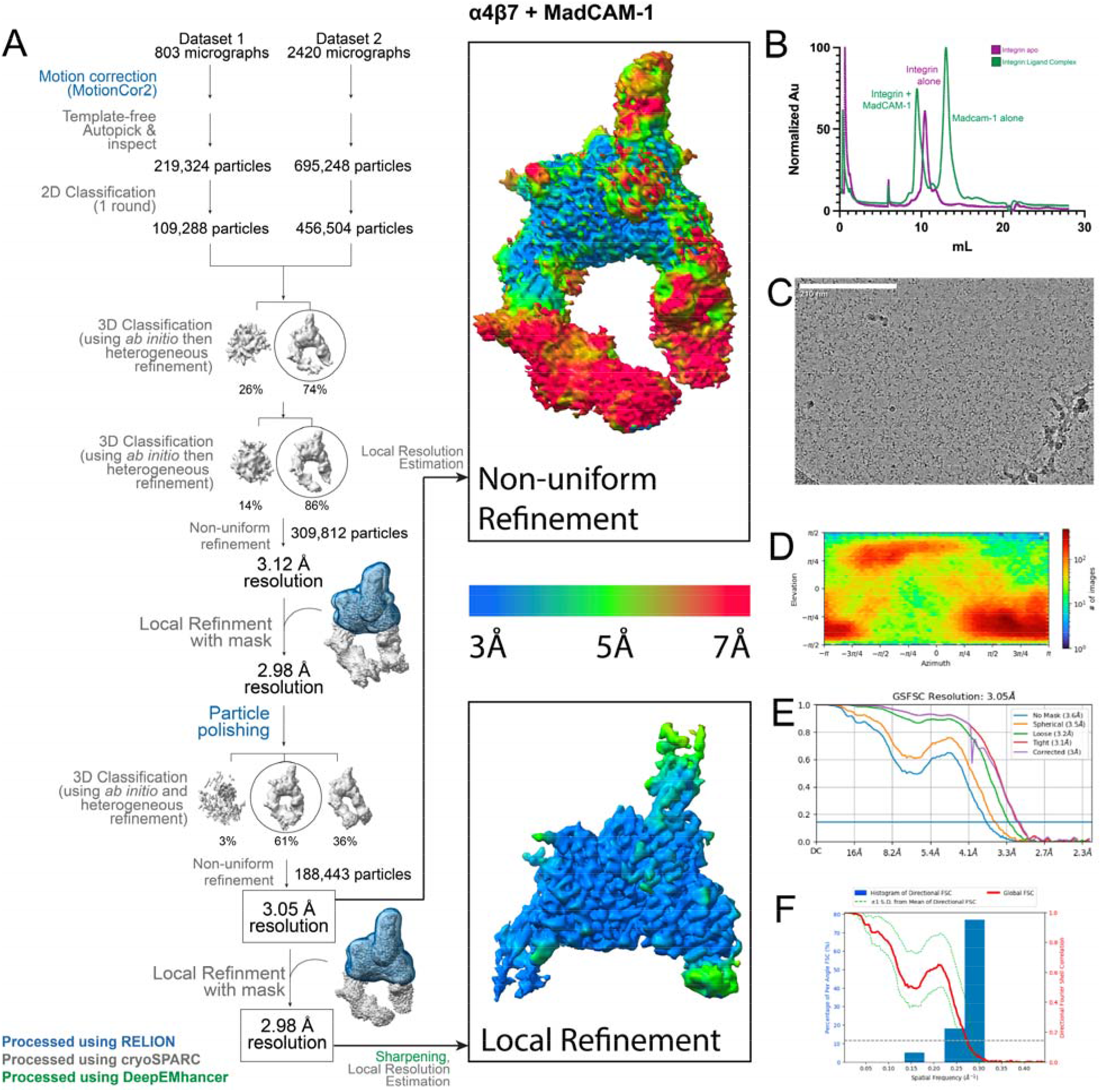
Data processing schematic for the α4β7:MadCAM-1 complex. **A,** A flowchart for the data processing pipeline of α4β7:MadCAM-1. Both the global and local refinements were used for model building. **B,** Size exclusion chromatography traces showing peak shift for ligand-bound integrin. **C,** Representative micrograph with 210nm scale bar, **D,** orientational distribution plot, **E,** gold-standard Fourier Shell Correlation (GSFSC) plot, and **F,** three-dimensional Fourier Shell Correlation (3DFSC) plot for the globally refined map.

**Supplemental Figure 3.**
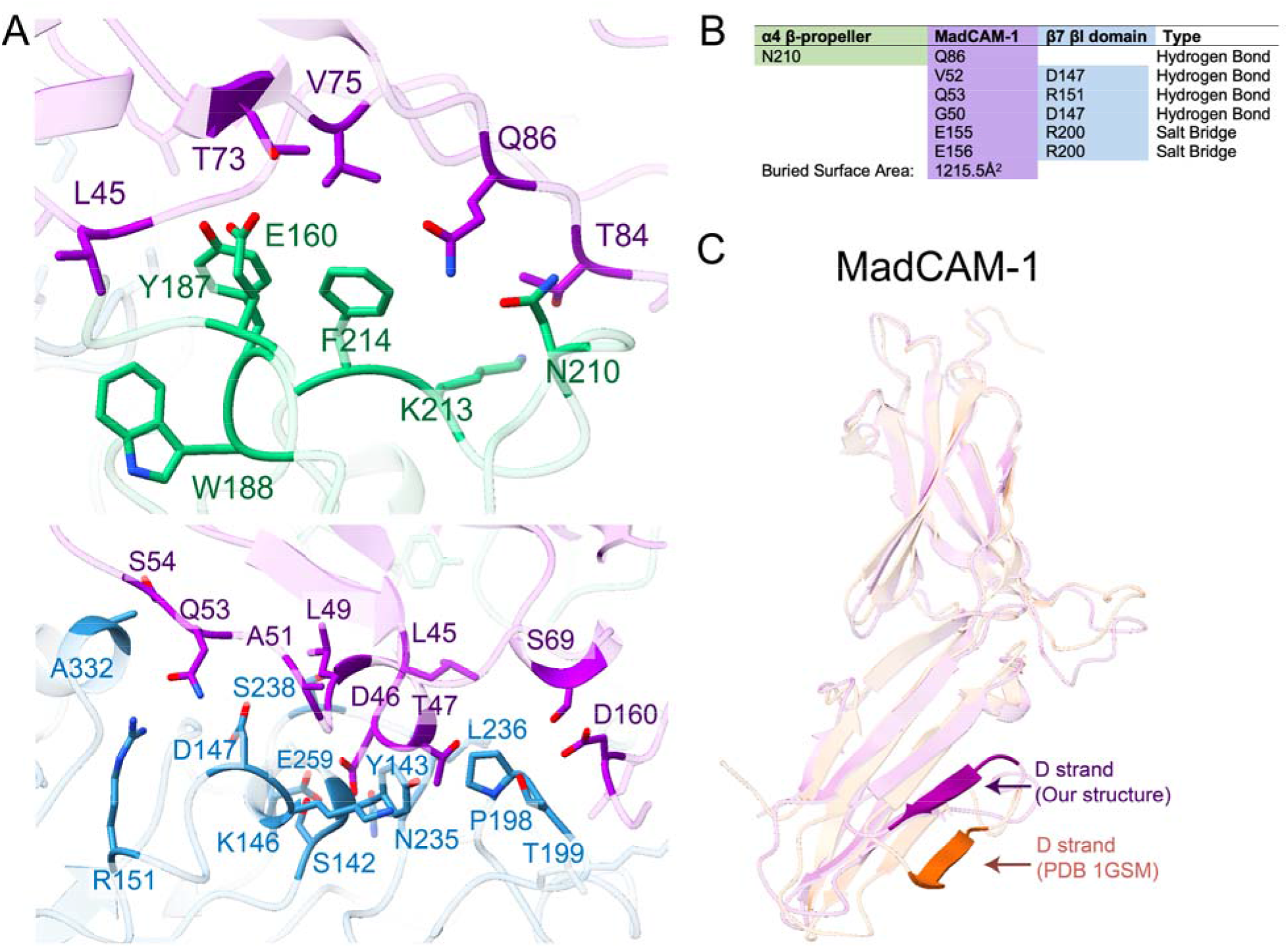
The molecular interface between integrin α4β7 and MadCAM-1. **A,** MadCAM-1 (purple) binding within the α4β7 groove is stabilized by contacts with both α4 (green, top) and more extensively with β7 (blue, bottom). **B,** The PISA server^57^ was used to determine electrostatic interactions in the α4β7:MadCAM-1 complex model **C,** MadCAM-1 undergoes a conformational shift upon binding to α4β7. The D strand in the first Ig-like domain of MadCAM-1 is shown in high opacity for both our structure (purple) and the crystal structure (orange, PDB 1GSM^25^). The conformation of the D strand in the crystal structure of MadCAM-1 alone would sterically clash with our integrin density.

**Supplemental Figure 4.**
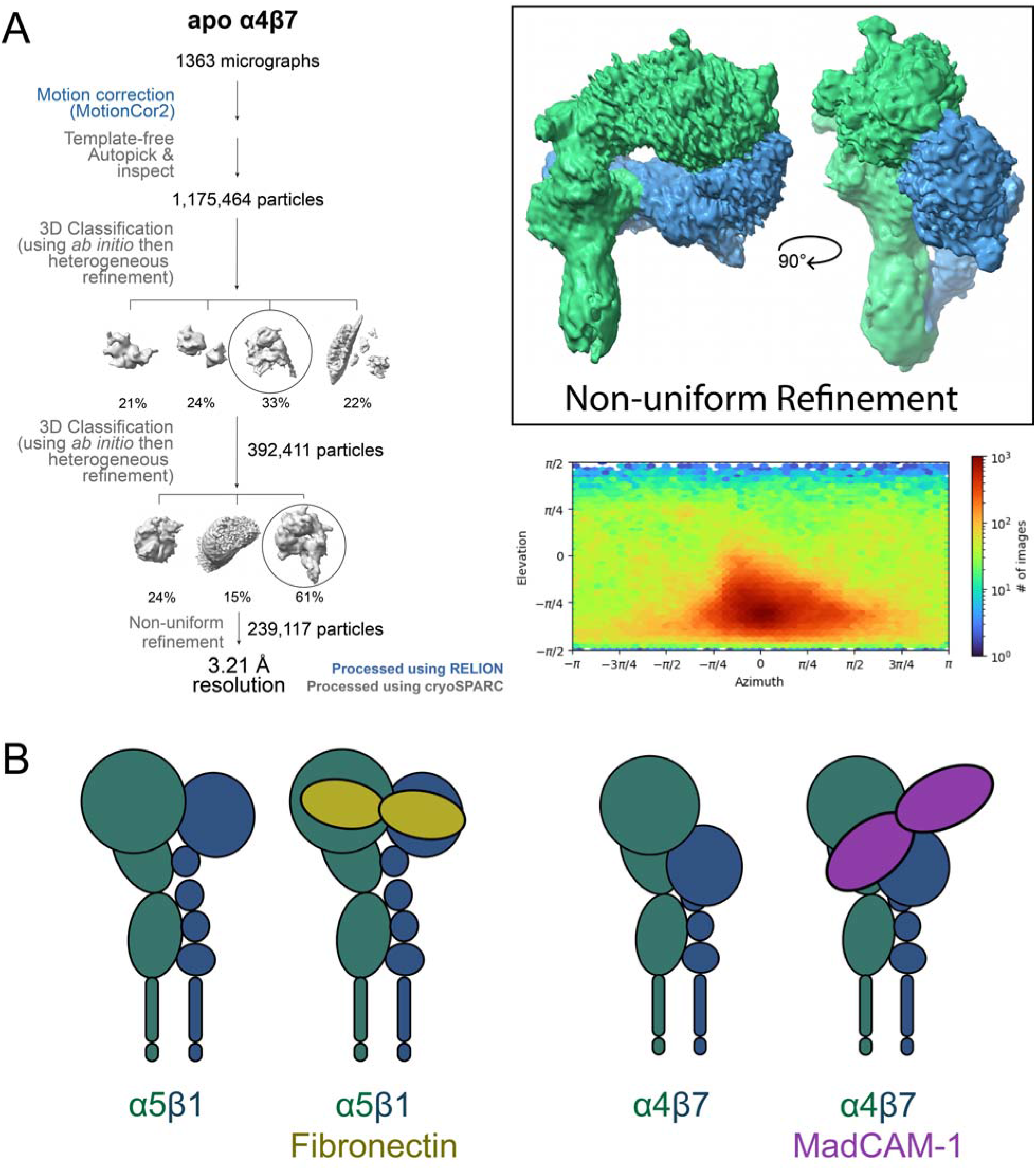
Integrin α4β7 has a noncanonical activation mechanism. **A,** A flowchart for the data processing pipeline of apo α4β7. A preferred orientation prevents us from building an atomic model to the generated density, although secondary structure is largely preserved. **B,** Unlike integrin α5β1, the compact half-bent state of α4β7 has a significant rotation (40°) at the headpiece. This may contribute to binding to MadCAM-1, which binds α4β7 perpendicular compared to how α5β1’s ligand fibronectin binds^31^.

**Supplemental Figure 5.**
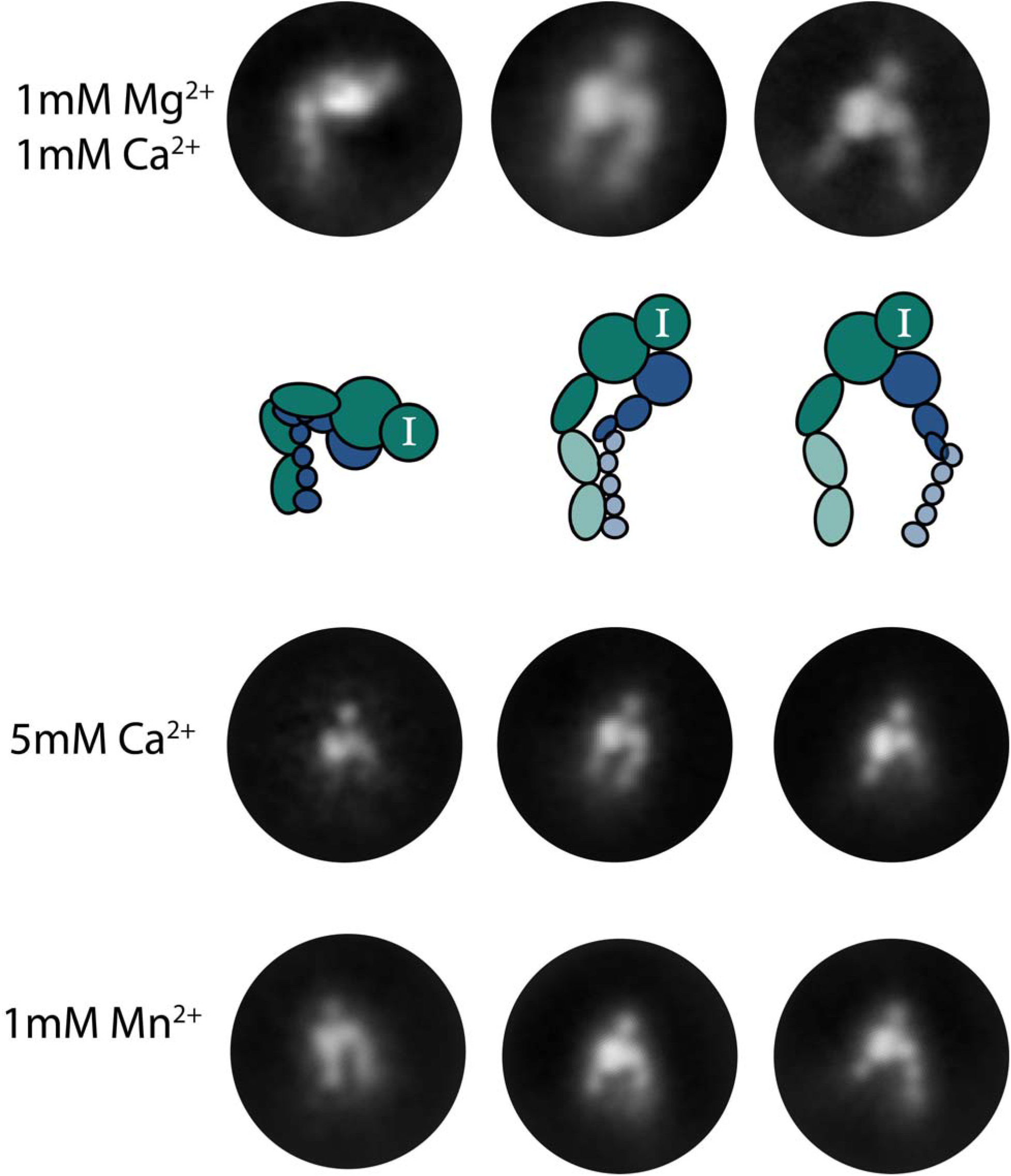
Integrin αEβ7 occupies canonical conformations. Negative stain electron microscopy (nsEM) 2D class averages of integrin αEβ7 in buffers of varying ions show that αEβ7 samples conformations similar to those previously described for other integrins, although unlike other leukocyte integrins αEβ7 appears to adopt a “half-bent” conformation. The I domain is clearly resolved in each class.

**Supplemental Figure 6.**
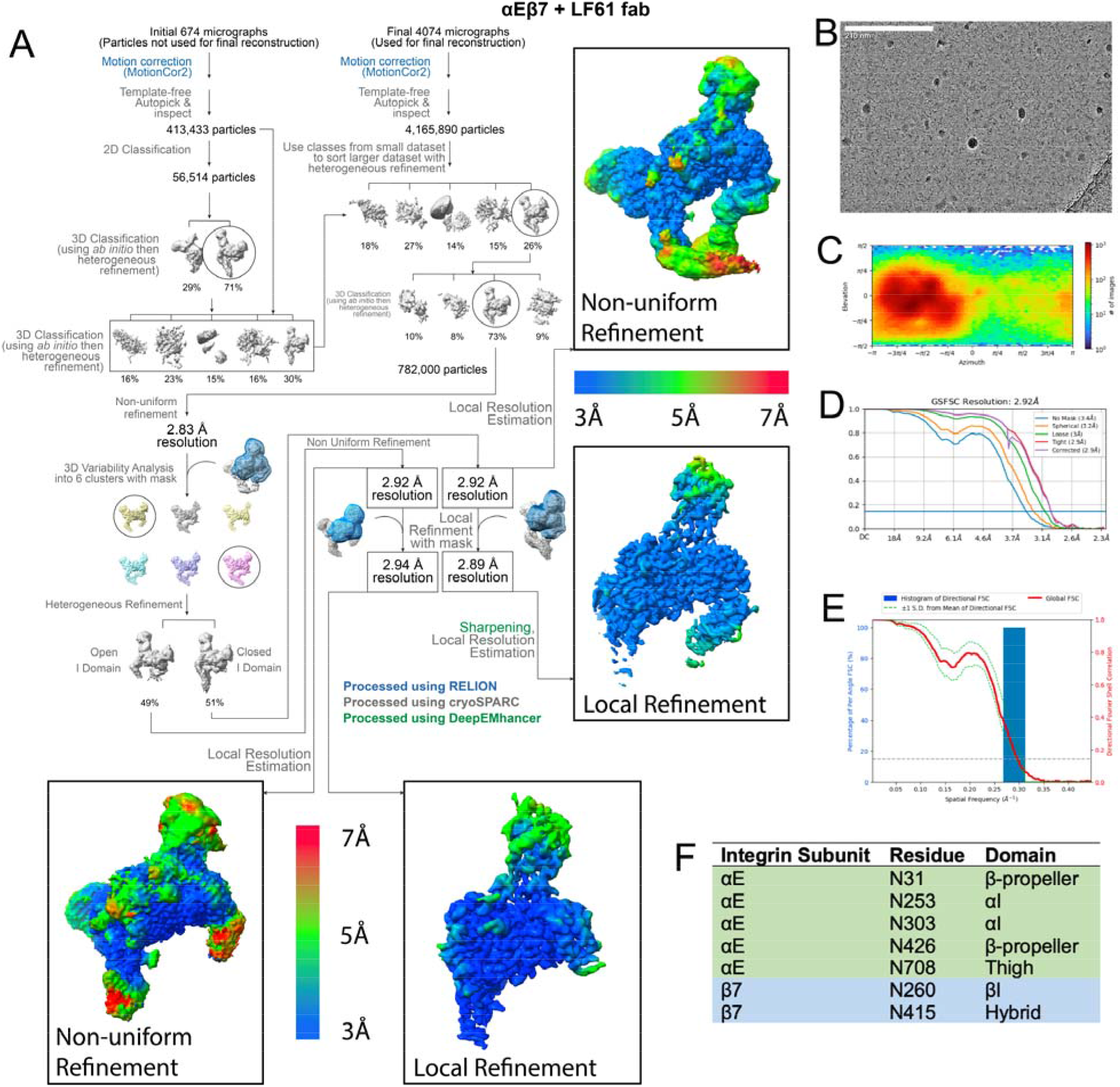
Data processing schematic for the αEβ7:LF61 Fab complex. **A,** A flowchart for the data processing pipeline of αEβ7:LF61 Fab. High-quality particles were sorted into two major classes; those that have an open I domain and those that have a closed I domain. The closed I domain structure, shown on the right with local resolution estimates, was used to model the inactive αEβ7 conformation. The open structure is shown below. **B,** Representative micrograph with 210nm scale bar, **C,** orientational distribution plot, **D,** gold-standard Fourier Shell Correlation (GSFSC) plot, and **E,** three-dimensional FSC (3DFSC) plot for the globally refined closed I domain map. **F,** Glycan residues and domains within the αEβ7:LF61 model.

**Supplemental Figure 7.**
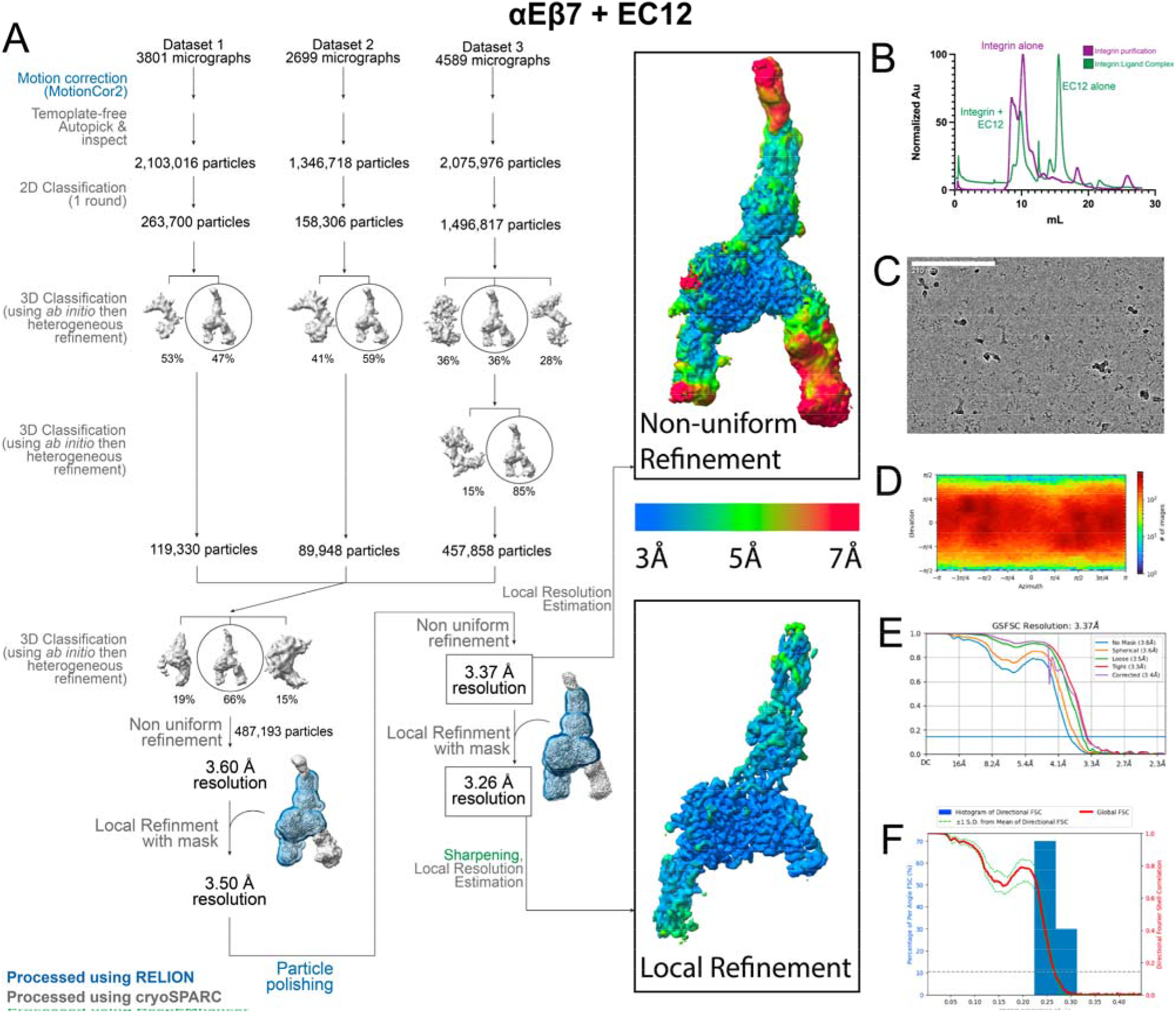
Data processing schematic for the αEβ7:EC12 complex. **A,** Flowchart for the data processing pipeline of αEβ7:EC12. Both the global and local refinements were used for model building. **B,** Size exclusion chromatography traces showing peak shift for ligand-bound integrin. **C,** Representative micrograph with 210nm scale bar, **D,** orientational distribution plot, **E,** gold-standard Fourier Shell Correlation (GSFSC) plot, and **F,** three-dimensional FSC (3DFSC) plot the globally refined map.

**Supplemental Figure 8.**
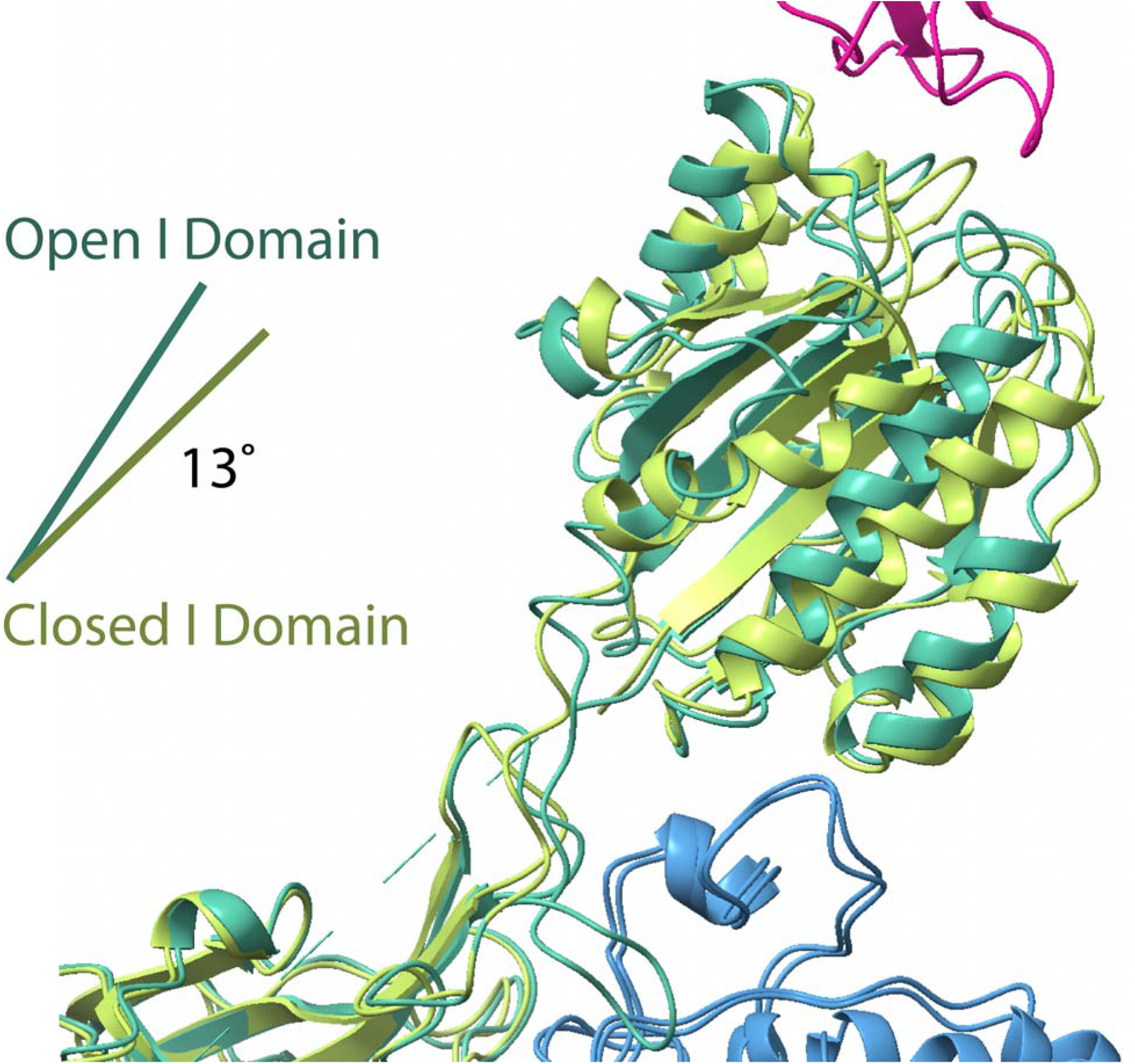
Structural differences between the apo closed and ligand-bound, open αE I domain. Despite extensive structural changes the location of the I domain relative to the rest of the integrin molecule remains the same when ligand-bound. The I domain has a ∼13° shift upon binding to E-cadherin. Models are aligned on the αE beta-propeller. Open I domain shown in spearmint, closed I domain shown in lime.

**Supplemental Figure 9.**
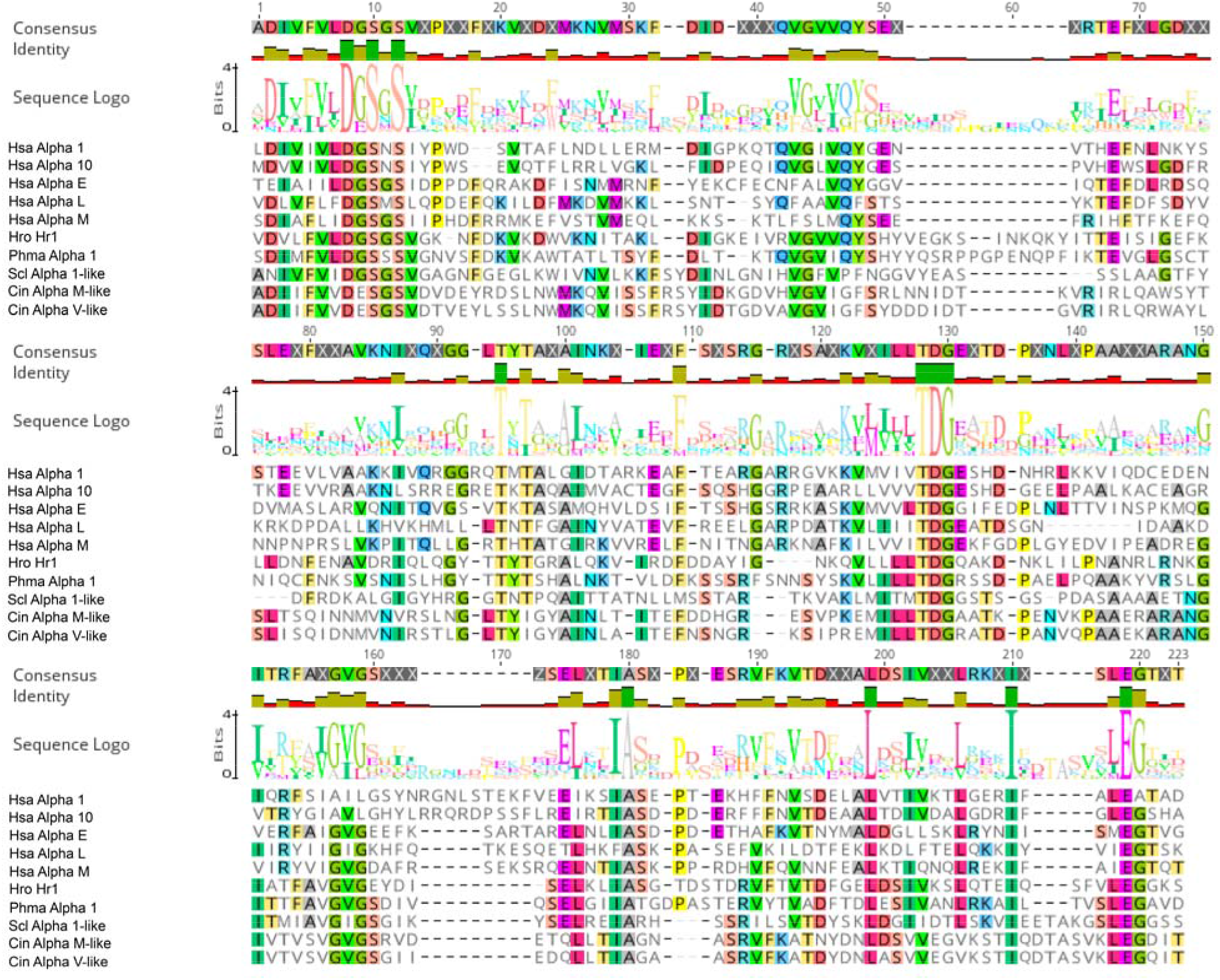
An alignment of representative integrin I domains used for Hidden Markov Model (HMM)-searching. A MUSCLE alignment of integrin I domains across Olfactores was generated in Geneious. Names and accession numbers are as follows: Hsa (*Homo sapiens*) Alpha 1 (NP_852478.1), Alpha 10 (AAC31952.1), Alpha E (EAW90480.1), Alpha L (NP_002200.2), Alpha M (AAH96346.1), Hro (*Halocynthia roretzi*) Hr1 (BAB21479.1), Phma (*Phallusia mammillata*) Alpha 1 (CAB3257102.1), Scl (*Styela clava*) Alpha 1-like (XP_039274816.1), Cin (*Ciona intestinalis*) Alpha M-like (XP_026691356.1), Alpha V-like (XP_026691784.1).

**Supplemental Figure 10.**
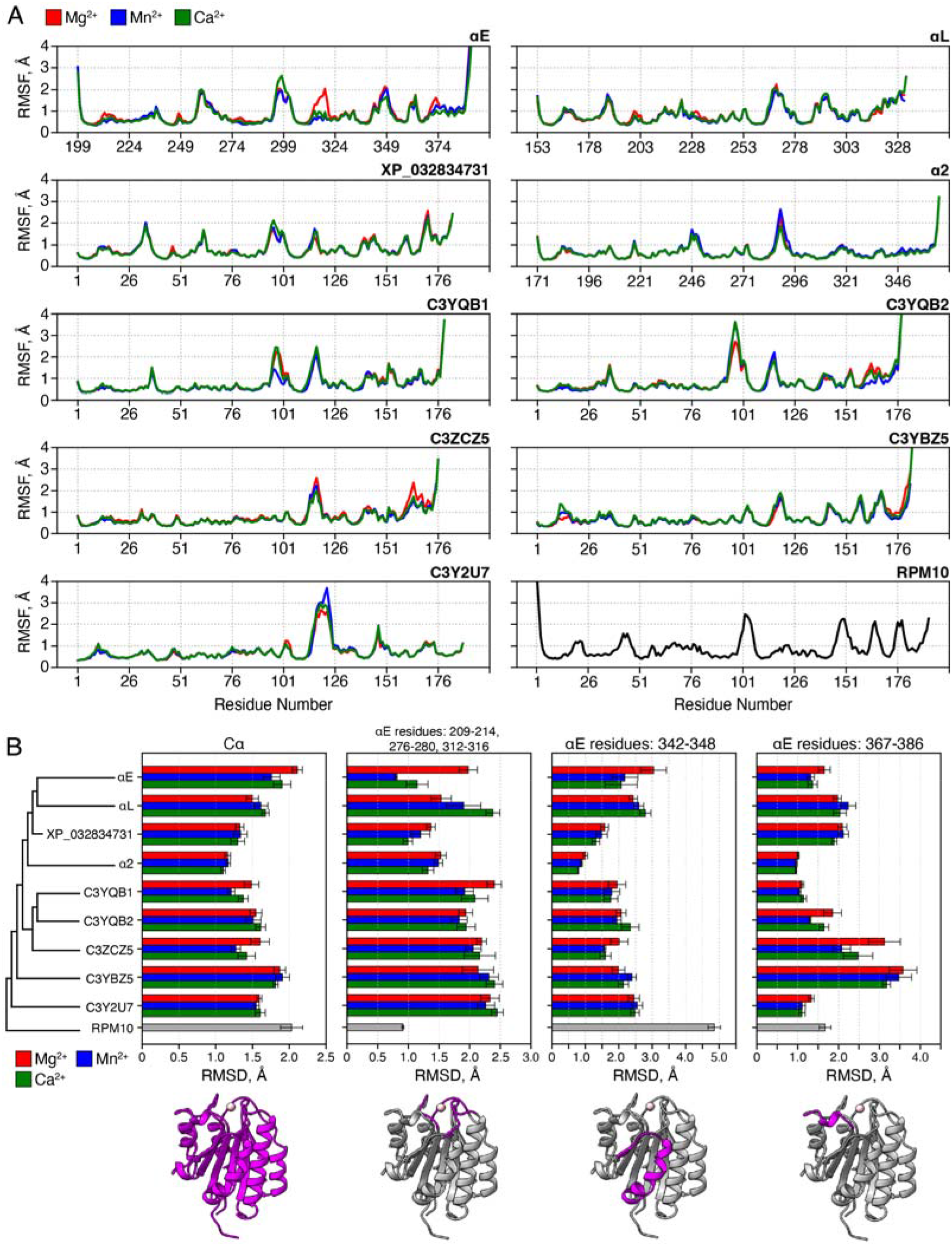
Molecular dynamics simulations reveal conserved dynamics in vWFA domains. **A,** Root mean square fluctuation (RMSF) of 10 simulated vWFA proteins with either Mg^2+^ (red), Mn^2+^ (blue), or Ca^2+^ (green) bound in the MIDAS site. For RPN10, no ion was bound. **B,** RMSD calculation of the simulated vWFA proteins. Color represents the cation bound in the MIDAS site (Mg^2+^, red; Mn^2+^, blue; Ca^2+^, green). Regions calculated, from left to right, all Cα atoms, ion coordinated loops, and sites of conformational change and colored magenta in the structure. Error bars represent standard error. For both (A) and (B), values are averaged across 10 independent replicates of 2.4µs long with only the last 2.0µs used for analysis and the initial simulation structure used as the reference.

**Supplemental Figure 11.**
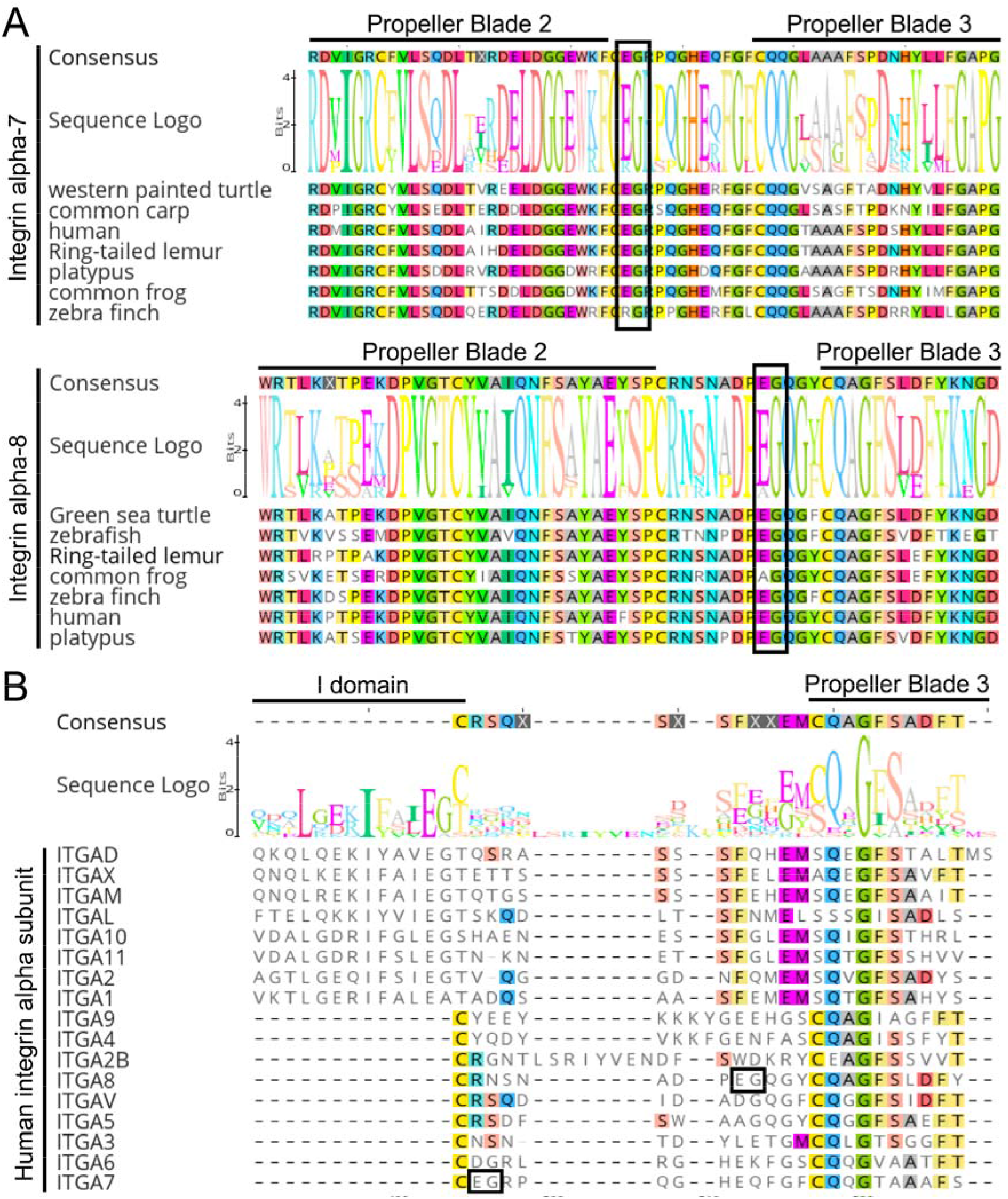
Extant integrins contain internal ligand-like motifs within the beta propeller. **A,** The integrin I domain was inserted between the second and third beta propeller domains of an ancestral integrin. We could not find conserved “I/LEGT” motifs in our HMM search, leaving the possibility open that the I/LEGT motif was already present in the ancestral integrin. Extant integrin subunits α7 (top) and α8 (bottom) contain EG motifs that are conserved within the subunit across vertebrates. **B,** An alignment of all human integrin subunits shows these EG motifs are not broadly conserved between integrins, leaving the origin of the internal ligand I/LEGT motif ambiguous. All alignments are MUSCLE alignments generated using Geneious.

**Supplemental Table 1.** Collection, refinement and validation statistics for cryoEM density maps and models. Collection details for all cryoEM maps presented in the text are shown above, and validation statistics for the models subsequently generated from those maps are shown below.

| | αEβ7:LF61<br>Closed I domain<br>(EMDB-xxxx)<br>(PDB xxxx) | αEβ7:LF61<br>Open I domain<br>(EMDB-xxxx)<br>(PDB xxxx) | αEβ7:EC12<br>(EMDB-xxxx)<br>(PDB xxxx) | α4β7:MadCAM-1<br>(EMDB-xxxx)<br>(PDB xxxx) | apo α4β7<br>(EMDB-xxxx) |
| --- | --- | --- | --- | --- | --- |
| **Data collection and processing** |  |  |  |  |  |
| Magnification | 36000x | 36000x | 36000x | 36000x | 36000x |
| Voltage (kV) | 200 | 200 | 200 | 200 | 200 |
| Electron exposure (e-/$\text{\AA}^2$) | 50 | 50 | 50 | 50 | 50 |
| Defocus range ($\mu\text{m}$, nominal) | 1.2-1.8 | 1.2-1.8 | 1.2-1.8 | 1.2-1.8 | 1.2-1.8 |
| Pixel size ($\text{\AA}$) | 1.122 | 1.122 | 1.122 | 1.122 | 1.122 |
| Symmetry imposed | C1 | C1 | C1 | C1 | C1 |
| Initial particle images (no.) | 4,165,890 | 4,165,890 | 5,307,702 | 914,572 | 1,175,464 |
| Final particle images (no.) | 402,099 | 379,901 | 487,193 | 188,443 | 239,117 |
| Map resolution ($\text{\AA}$) | 2.92 | 2.93 | 3.37 | 3.05 | 3.21 |
| FSC threshold | 0.143 | 0.143 | 0.143 | 0.143 | 0.143 |
| Map resolution range ($\text{\AA}$) | 2.5-10.2 | 2.5-16.8 | 2.8-10.9 | 2.5-46.5 | 2.7-47.6 |
| **Refinement** |  |  |  |  |  |
| Initial model used (AlphaFold or PDB code) | AlphaFold<br>PDB: 3V4P |  | AlphaFold<br>PDB: 7NLW, 4ZT1 | AlphaFold-multimer |  |
| Model resolution ($\text{\AA}$) | 2.92 | | 3.37 | 3.05 | |
| FSC threshold | 0.143 |  | 0.143 | 0.143 |  |
| Model resolution range ($\text{\AA}$) | 2.5-10.2 | | 2.8-10.9 | 2.5-46.5 | |
| Model composition |  |  |  |  |  |
| Non-hydrogen atoms | 8933<br>1135 |  | 9792<br>1250 | 9150<br>1162 |  |
| Protein residues | 22 |  | 18 | 22 |  |
| Carbohydrates |  |  |  |  |  |
| B factors ($\text{\AA}^2$) | | | | | |
| Protein | 113.5 |  | 119.7 | 87.1 |  |
| Ligand | N/A |  | N/A | N/A |  |
| R.m.s. deviations |  |  |  |  |  |
| Bond lengths ($\text{\AA}$) | 0.012 (11) | | 0.012 (12) | 0.012 (10) | |
| Bond angles ($^\circ$) | 1.949 (52) | | 2.038 (61) | 2.048 (73) | |
| Validation |  |  |  |  |  |
| MolProbity score | 0.78 |  | 0.82 | 0.97 |  |
| Clashscore | 0.17 |  | 0.05 | 0.22 |  |
| Poor rotamers (%) | 0.21 |  | 0.09 | 0.20 |  |
| Ramachandran plot |  |  |  |  |  |
| Favored (%) | 96.63 |  | 95.73 | 94.37 |  |
| Allowed (%) | 3.19 |  | 3.95 | 5.55 |  |
| Disallowed (%) | 0.18 |  | 0.32 | 0.09 |  |

## Supplemental Information Legends

**Supplemental Video 1**. Integrin α4β7 is flexible when bound to MadCAM-1. 3DFlex analysis was used to analyze continuous movement within the α4β7:MadCAM-1 complex. The β7 subunit (blue) shows hinged motion at the hybrid domain, and there is coordinated rotational movement between the α4 (green) and β7 subunits at the lower leg.

**Supplemental Video 2**. Compact αEβ7 stochastically samples an internally-liganded state. 3D Variability analysis was used to separate compact αEβ7:LF61 particles into states with or without the internal ligand engaged. αE is represented in green and β7 in blue. The internal ligand is the central density with high variability between frames.

**Supplemental Video 3**. Integrin αEβ7 is flexible when bound to E-Cadherin. 3DFlex analysis was used to analyze continuous movement within the αEβ7:EC12 complex. The active β7 subunit (blue) shows a similar hybrid domain motion as in α4β7:MadCAM-1. There is also some flexible motion between E-cadherin domains EC1 and EC2 (pink). αE is represented in green.

**Supplemental Data 1**. Alignment of human β-integrin protein sequences. The alignment used to generate the β-integrin subunit tree presented in Figure 2 in PHYLIP format. Names are presented as NCBI accession numbers.

**Supplemental Data 2**. Alignment of human α-integrin protein sequences. The alignment used to find the N- and C-terminal I domain insertion regions presented in Figures 6A and Supplemental Figure 11 in PHYLIP format. Names are presented as gene names.

**Supplemental Data 3**. Alignment of *Olfactores* I domain protein sequences. The alignment used to generate the Hidden Markov Model (HMM) used to identify candidate cephalochordate I domain-like proteins in FASTA format. Names for human genes are given “Alpha*,” while all other names are presented as NCBI accession numbers.

**Supplemental Data 4**. Alignment of I domains and cephalochordate outgroup vWFA domains. The alignment used to generate the phylogenetic tree presented in Figure 6C. Names are presented as NCBI accession numbers for integrin I domains or Uniprot identifiers for cephalochordate vWFA domains.

